# Distinct and Combined Interferon-ɑ/β-receptor-1 Loss in Neurons and Astrocytes Disrupt Brain Energy Metabolism and Drive Parkinsonian Dementia

**DOI:** 10.1101/2025.08.14.670043

**Authors:** Erika B. Villanueva, Zala Zebec, Jon Lundstrøm, Jens V. Andersen, Filippa L. Qvist, Emil W. Westi, Andrea Marin, Gisela Jimenez-Duran, Lluís Riera-Ponsati, Emilie Tresse, Oliver Kretz, Desiree Loreth, Tobias Goldmann, Thomas Blank, Marco Prinz, Blanca I. Aldana, Matthias Mann, Niels H. Skotte, Shohreh Issazadeh-Navikas

## Abstract

Dysregulated interferon-alpha/beta-receptor 1 (IFNAR1) signaling was recently identified to contribute to the development of sporadic Parkinson’s Disease (PD) into PD with Dementia (PDD). The molecular, cellular, and phenotypic impacts of brain IFNAR1 loss in aging have not been explored *in vivo*, which may reveal novel disease mechanisms and therapeutic targets. Here it is shown that baseline IFNAR1 expression varies in the major brain cell types, including neurons and astrocytes, and is differentially affected in PD and Lewy Body Dementia patients compared to unaffected controls. Neuron- and astrocyte-specific transcriptomic and proteomic alterations in *Ifnar1^−/−^* mice implicate mitochondrial defects and synergistic dysfunctional neurotransmission upon IFNAR1 loss, leading to glucose hypermetabolism measured by functional metabolic analysis. Consequently, *Ifnar1^−/−^* mice exhibited PDD-like pathogenesis, including dopaminergic cell loss in the substantia nigra, cortical neurodegeneration, Lewy-body-like inclusions, neuroinflammation, and progressive PDD-like behavior deficits. Brain cell-specific IFNAR1 loss examined *in vivo* revealed delayed but distinct development of PDD-like phenotypes, where neuropathology, motor, and cognitive behavior deficits were specifically recapitulated only in mice lacking neuronal IFNAR1, and behavior resembling neuropsychiatric abnormalities recapitulated only in mice lacking astrocytic IFNAR1. This work supports a crucial role of IFNAR1 in brain homeostasis and emphasizes a need for understanding neurodegenerative pathophysiology in cell-specific contexts.

**Highlights:**

- *IFNAR1* and related type-I IFN genes are differentially expressed among major brain cell types in Parkinson’s Disease, Lewy Body Dementia, and unaffected controls
- Early molecular alterations in *Ifnar1^−/−^* mice show lack of immunomodulation contributing to neuroinflammation, mitochondrial defects, and dysregulated energy metabolism
- *Ifnar1^−/−^* mice develop a progressive Parkinsonian-like disease phenotype, including dopaminergic cell loss in substantia nigra, cortical neurodegeneration, phosphorylated (p)alpha-synuclein^+^ and pTau^+^ Lewy-body-like inclusions, neuroinflammation, and progressive motor, cognitive, and neuropsychiatric disturbance-like behavior deficits
- Neuropathologies, motor, and cognitive deficits are recapitulated in mice lacking neuronal IFNAR1 (Syn1^Cre^;*Ifnar1^fl/fl^*) whereas neuropsychiatric abnormalities are recapitulated in mice lacking astrocytic IFNAR1 (GFAP^Cre^;*Ifnar1^fl/fl^*)

## Introduction

The molecular mechanisms underlying dementia formation in sporadic Parkinson’s disease (PD), which can develop more than 10 years after motor symptom onset, are not well understood (1). A large percentage of PD with dementia (PDD) cases have neuropathological similarities to other dementias such as Lewy Body Dementia (LBD) and Alzheimer’s disease (AD), including neuroinflammation and gliosis associated with phosphorylated-tau (pTau) and amyloid-beta (Aβ) proteinopathies, the latter correlating with severity of dementia development (1–4). Brain energy metabolism disturbances have been reported in Parkinsonian disorders (5–8) and AD (9), with region-specific alterations in glucose metabolism corresponding with severity of both motor symptoms (7, 8) and dementia (5). As metabolic alterations have been observed prior to dementia onset in both PD (6) and AD (9), understanding the complex connection between neuroinflammation, energy metabolism, proteinopathy, neurodegeneration, and clinical outcomes is imperative for designing disease-modifying therapeutics.

Dysregulated type-I interferon (IFN) signaling, specifically the interaction between the interferon-beta (IFNβ) cytokine and the interferon alpha/beta receptor (IFNAR)1, has been implicated in the conversion of human sporadic PD to PDD (10), and we have investigated its role in brain homeostasis through a series of studies. Dysfunctional type-I IFN in PD patients and lack of IFNβ*-*IFNAR signaling in mice causes extrusion of damaged mitochondrial DNA, which was found to be sufficient to initiate and propagate PDD-like neuropathology upon injection into healthy brains (11). Lack of IFNβ in mice (*Ifnb^−/−^*) was sufficient to induce spontaneous and progressive behavioral abnormalities and neuropathologies resembling PDD (12, 13). Mice lacking IFNAR1 (*Ifnar1^−/−^*) in general or specifically in Nestin^+^ neural ectodermal cells (Nes^Cre^;*Ifnar1^fl/fl^*) also develop alpha-synuclein (α-syn)^+^ Lewy body (LB)-like pathology upon aging (13), implicating loss of neuronal and/or glial IFNAR1 signaling in progressive neurodegeneration. An independent study found that young *Ifnb^−/−^* and *Ifnar1^−/−^*mice have compromised synaptic plasticity due to impaired astrocytic glutamate transporter function (14); however, specific neuronal versus astrocytic dysfunction in *Ifnar1^−/−^* mice upon aging has not been explored. Identifying early brain cell-specific molecular alterations upon IFNAR1 loss *in vivo* may aid in understanding pathogenesis resembling PDD.

Here, publicly available single nuclei RNA sequencing (snRNA-seq) datasets were used to establish baseline expression of *IFNAR1* in major human brain cell types of healthy individuals, which were compared with samples from patients with LBD and PD. Investigating both human and mouse brains unveiled differential *IFNAR1* and related type-I IFN gene expression patterns in neurons, astrocytes and other brain cell types which were further distinguished in LBD and PD, suggesting unique requirements in brain homeostasis and potentially distinct impacts on neurodegenerative disease progression and dementia development. Next, cell-specific molecular differences were identified in *Ifnar1^−/−^* versus wild-type (Wt) mouse brains using an integrative approach, combining global and unbiased snRNA-seq, liquid chromatography tandem mass spectrometry (LC-MS/MS)-based proteomics, and functional metabolic ^13^C isotope tracing, which together revealed mitochondrial dysfunction and disturbed brain energy metabolism upon IFNAR1 loss. Consequently, *Ifnar1^−/−^* mice developed PDD-like neuropathology, neuroinflammation, and progressive motor, neuropsychiatric, and cognitive behavior deficits. Furthermore, cell-specific IFNAR1 loss in neurons (Syn1^Cre^;*Ifnar1^fl/fl^* mice) or astrocytes (GFAP^Cre^;*Ifnar1^fl/fl^* mice) developed distinct aspects of Parkinsonian disease-like phenotypes observed in *Ifnar1^−/−^* mice: neuronal IFNAR1 loss was sufficient to induce to PDD-like neuropathologies and subsequent motor and cognitive dysfunctions, whereas astrocytic IFNAR1 loss led to specific manifestation of behavior resembling neuropsychiatric abnormalities. These results support a pivotal role of IFNAR1 in brain homeostasis, which can lead to distinct pathological outcomes depending on brain cell type dysfunction, while their concerted dysregulations are required for development of full-blown PDD-like disease features. Our findings highlight a need for understanding neurodegenerative pathophysiology in cell-specific contexts. Concerted and targeted restoration of dysfunctional Type-I-IFN signaling through IFNAR1 in neurons and astrocytes may therefore mitigate development of dementia resembling PDD.

## Materials and Methods

### Human single nuclei (sn)RNA-seq data analysis

The RNA single cell type data released by the Allen Brain Institute (https://portal.brain-map.org/atlases-and-data/rnaseq/human-m1-10x) was used to obtain IFNAR1 expression values (nTPM) in specific brain cell subsets, including excitatory and inhibitory neurons, astrocytes, Oligodendrocyte Precursor Cells (OPCs), oligodendrocytes and microglia (15). To compare IFNAR1 and related Type-1 IFN gene expression between unaffected controls and Lewy Body dementia (LBD) or Parkinsońs disease (PD) patients within specific brain cell subsets, data from the publicly accessible snRNA-seq study ((16), available at: https://singlecell.broadinstitute.org/single_cell/study/SCP1768/) were used. Scaled mean expression values of IFNAR1 from each individual donor ID were extracted for human dopaminergic (DA) neurons, non-DA neurons, astrocytes, OPCs, oligodendrocytes and microglia cluster datasets.

### Mice

Mice were housed in standard facilities (13). *Ifnb^−/−^*(13) and *Ifnar1^−/−^*(17) mice were backcrossed more than 20 generations on to a C57BL/6J background. Wild-type (Wt) mice were *Ifnb^+/+^*and *Ifnar1^+/+^* mice on the C57BL/6J background. C57BL/6J mice carrying loxP-flanked *Ifnar1* (*Ifnar1^fl/fl^*) (18) were crossed with C57BL/6J transgenic mice expressing Cre recombinase under neuron-specific synapsin-1 promoter (Syn1^Cre^) (19) or astrocyte-specific glial fibrillary acidic protein promotor (GFAP^Cre^) (20) to obtain Syn1^Cre^;*Ifnar1^fl/fl^* and GFAP^Cre^;*Ifnar1^fl/fl^* mice, respectively. Behavior cohort numbers and sample sizes for biochemical analyses were determined as previously described (12). Males and females of roughly equal proportions were used for snRNA-seq, flow cytometry, qPCR, behavior, and immunohistochemistry. As higher disease prevalence and severity occurs in males in human PD (21), male behavior-tested mice were used for further evaluations, including proteomics, and metabolic studies, electron microscopy, and immunoblots. Five familial AD mutation-containing (5xFAD) transgenic mice (22), which develop significant extracellular Aβ pathology, were used for Aβ antibody verification.

### Brain tissue preparation

For snap-frozen tissues, mice were sacrificed by cervical dislocation. Brains were removed swiftly and placed on a petri dish over ice and microdissected for cortex, hippocampus, and basal ganglia. Tissues were snap-frozen in liquid nitrogen and kept at -80°C until use for snRNA-seq, LC-MS/MS, or immunoblots. A summary of the brain regions investigated in this study, their relevance to known pathology in PD and PDD, investigation methods used, and the age of mice at the time of investigation, is shown in Table 1.

**Table 1.**
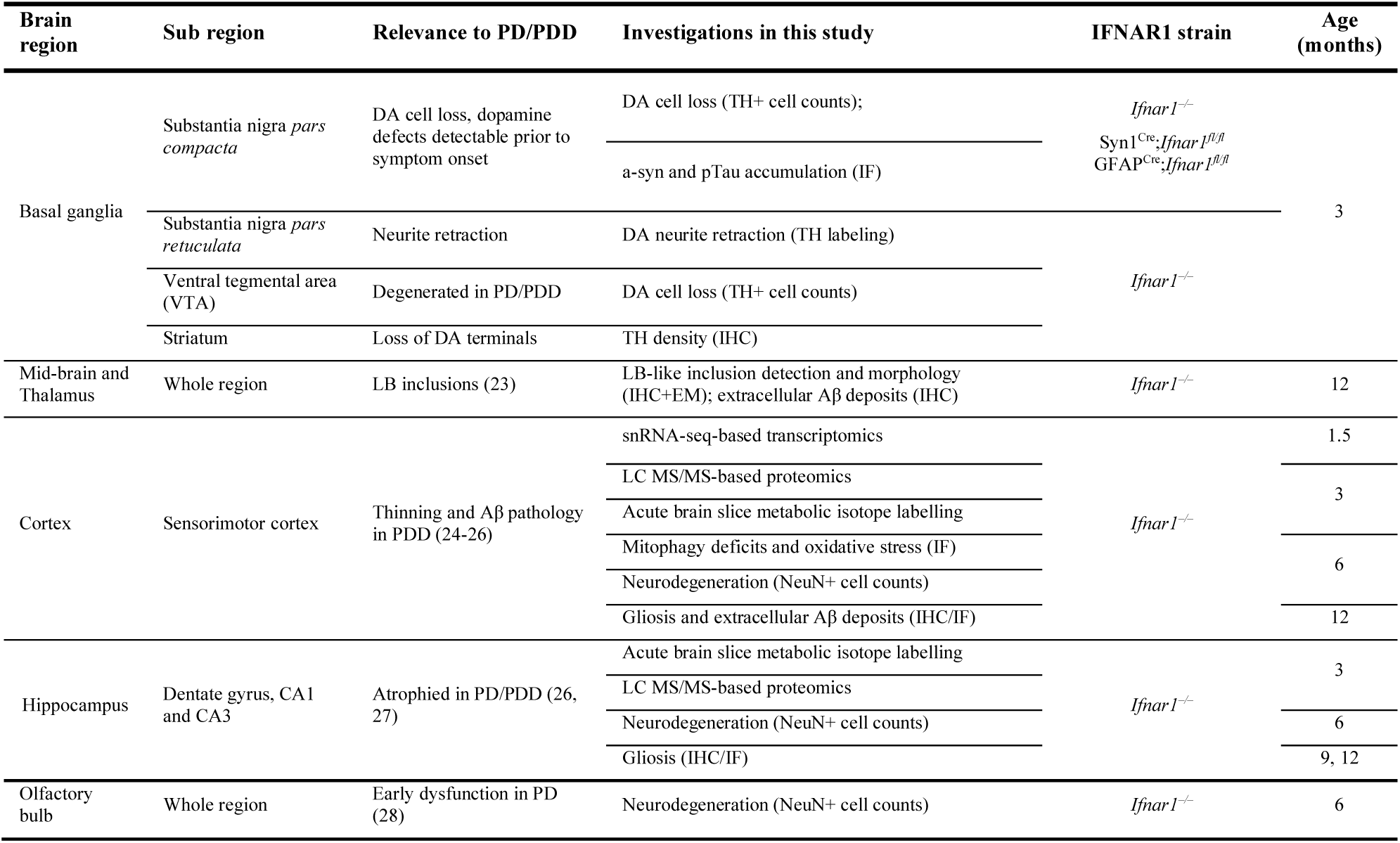
PD/PDD-like neuropathologies by brain region and their investigation in this study.

For paraffin-embedded tissues, mice were placed under isofluorane-induced anesthesia and intracardially perfused for 1 min with phosphate-buffered saline (PBS), followed by freshly prepared aqueous PBS-buffered 4% paraformaldehyde (PFA, Sigma, P6148) fixative solution. Brains were removed and kept in the same fixative at 4°C for 18 h before alcohol dehydration and paraffin embedding. A microtome (Thermo Scientific, HM355S) was used to prepare 6 µm thick serial coronal sections onto glass slides (Thermo Scientific, 10149870).

### Mouse snRNA-seq

#### Nuclear isolation

Extraction of nuclei from snap-frozen cortical tissue was performed using a modified version of previously described protocols(29, 30). Briefly, nuclei isolation media (NIM) (250 mM sucrose, 25 mM KCl, 5 mM MgCl_2_, and 10 mM tris buffer, pH 8) was prepared. Each cortex was transferred to a precooled Dounce homogenizer containing 1 mL of ice-cold homogenization buffer (NIM supplemented with 1 mM DTT (Invitrogen), 0.4 U/μL RNase inhibitor (Takara), 0.2 U/μL Suprasin (Invitrogen), and 0.1% v/v Triton X-100) and dissociated on ice using five strokes of the loose pestle, followed by fifteen strokes of the tight pestle. Homogenates were filtered through 40 μm strainers and nuclei were collected by centrifugation (1000 *g*, 10 min at 4 °C). Pellets were gently resuspended in 250 μL homogenization buffer each and mixed with 250 μL 50% iodixanol (50% vol/vol iodixanol, 150 mM KCl, 30 mM MgCl_2_, 60 mM tris buffer, pH 8, 0.4 U/μL RNase inhibitor, 0.2 U/μL Suprasin, and 1 mM DTT (Invitrogen)). 500 μL 29% iodixanol (29% vol/vol iodixanol, 150 mM KCl, 30 mM MgCl_2_, 60 mM tris buffer, pH 8, 0.4 U/μL RNase inhibitor, 0.2 U/μL Suprasin, and 1 mM DTT) was added to a 1 mL Beckman tube, precoated with 0.5% BSA in PBS with RNase inhibitor, and 500 μL of the nuclei suspension was slowly layered on top. Samples were spun at 14,000 g for 22 min at 4 °C in a TLS 55 rotor (MAX-XP ultracentrifuge, Accel 3, Decel 2). Supernatants were removed, and nuclei were resuspended in 0.5% BSA in phosphate-buffered saline (PBS) with RNase inhibitor and filtered through a 35 μm strainer. Nuclei counts were determined by trypan blue staining and hemocytometer count.

#### 10x Chromium sequencing

16,000 nuclei per sample were loaded on the 10x Chromium chip for library preparation using the Single Cell 3’ v3 chemistry according to manufacturer instructions. For cDNA amplification, 12 PCR cycles were applied. For each sample, 230 ng of purified cDNA was used for library construction. The libraries were diluted to a concentration of 10 nM and pooled for sequencing. After denaturation, the library pool was diluted to a loading concentration of 300 pM and sequenced on a Novaseq6000 in paired-end mode (2×100) (read 1:28 cycles, read 2:91 cycles, and i7 index: 8 cycles) at an average depth of 50,000 reads per nucleus.

#### Processing and quantification of 10x Chromium single-nuclei transcriptomes

Raw data were demultiplexed, aligned, and quantified using Cell Ranger version (v)3.1.0. To allow for counting of unspliced pre-mRNA, a custom version of the mm10-3.0.0 mouse reference genome provided by 10x Genomics was used, where the feature ‘type of transcripts’ was changed from ‘transcript’ to ‘exon’.

#### Single-nuclei data integration and analysis

Data analysis was performed in R using Seurat v3.1.5(31) and Conos v1.2.1(32). Briefly, the -barcode matrices were filtered to include cells meeting the following criteria: 200<nFeature_RNA<6000, nCount_RNA>1000, and percent_MT<10. Doublets were filtered out using the Scrublet package (33) and clusters containing mixed cell signatures were removed. Data were log-normalized, and individual samples were integrated using the fast mutual nearest neighbors (fastMNN) method(34) based on the top 3,000 variable features. Clustering and UMAP visualization were performed using the first 40 dimensions of the MNN reduction. The integrated dataset was annotated using canonical correlation analysis (CCA) based label transfer in Seurat with the Allen Brain Map Mouse Whole Cortex and Hippocampus SMART-seq dataset [https://portal.brain-map.org/atlases-and-data/rnaseq/mouse-whole-cortex-and-hippocampus-smart-seq] as the reference. Of note, very few astrocytes were detected in two samples (one Wt and one *Ifnar1^−/−^*) and were therefore excluded from astrocyte analysis to avoid technical overrepresentation of missing values. Differential gene expression analysis for each cortical cell type was performed using the getPerCellTypeDE() function in Conos. For pseudobulk data, differential gene expression analysis was performed using the getPerCellTypeDE() function in Conos, with all nuclei assigned to the same label for global pseudobulk comparison. Gene set enrichment analysis was performed using GSEA software (UC San Diego and Broad Institute)(35) and gene set collections from MsigDB(36). Total number of genes and DE genes per class were visualized using InteractiVenn(37). The upregulated KEGG pathway hits were visualized using *ggplot2,* for KEGG_PARKINSONS_DISEASE visualization the genes from GOBP_Oxidative_Phosphorylation were used to annotate mitochondria related genes.

### Liquid Chromatography Tandem Mass Spectrometry (LC-MS/MS)

#### Sample preparation for mass spectrometry

Snap-frozen cortex and hippocampus tissues were lysed in SDC alkylation and reduction lysis buffer (1% (w/v) sodium deoxycholate, 100 mM Tris pH 8, 40 mM CAA, 10 mM TCEP) using mechanical homogenizer (IKA® ULTRA-TURRAX® disperser, Merck) for 20 s. Lysates were boiled at 95°C for 10 min with continuous vortexing at 1200 rpm on a thermomixer (Eppendorf) to denature proteins. Samples were then sonicated using the Covaris Adaptive Focused Acoustics (AFA) sonication system (450W power, 50% duty factor, 200 cycles per burst) (Covaris, USA). Protein concentrations were measured using BCA assay (ThermoFisher Scientific). Proteins were digested overnight using trypsin (1:100 w/w, Sigma-Aldrich) and LysC (1/100 w/w, Wako) at 37°C. The next day fresh enzyme was added for another 2 h. The peptides were acidified by 1% trifluoroacetic acid (TFA) (Merck) to quench digestion. Peptide concentration was estimated using Nanodrop and peptide mixture was purified by solid phase extraction in Stage-Tips (SDB-RPS material, two 14-gauge plugs) for desalting and concentration. Peptides were washed twice with isopropanol/1% TFA and subsequently twice with 0.2% TFA, then eluted with 80% acetonitrile/1% ammonia. Eluted samples were reduced by vacuum centrifugation at 60°C and peptide concentration was determined by Nanodrop (ThermoFisher Scientific) measurement at A280 nm. Sample concentrations were adjusted to 200 ng per injection in buffer A* (5% ACN / 0.1% TFA) for LC-MS/MS analysis.

#### Preparation of mouse brain spectral library

High-pH reversed-phase fractionation was used to generate a deep precursor library for data-independent (DIA) MS analysis. Purified peptides were pooled from 6 brain regions, which were fractionated at pH 10 with the spider-fractionator (24 fractions from both cortex and hippocampus and 8 fractions from striatum, cerebellum, mid brain, and the olfactory bulb, respectively (80 fractions in total)(38). 6-18 μg of purified peptides were separated on a 30 cm C18 column in 100 min gradients and concatenated automatically into either 8 or 24 fractions with either 60- or 90-seconds exit valve switches for 24 or 8 fractions, respectively. Peptide fractions were vacuum-dried and reconstituted in buffer A* for LC-MS analysis.

#### LC-MS/MS Analysis

Samples were analyzed with nanoflow Easy-nLC 1200 (Thermo Fisher Scientific, Denmark) connected to a trapped ion mobility spectrometry quadrupole time-of-flight mass spectrometer (TimsTOF Pro, Bruker Daltonik GmbH, Germany) with a nano-electrospray ion source (Captive spray, Bruker Daltonik GmbH). Peptides were separated on a 50 cm in-house packed column (75 μm inner diameter × 50 cm length) with 1.9 μm ReproSilPur C18-AQ silica beads (Dr. Maisch, Germany). Column temperature was kept at 60°C using an in-house made column oven. Peptide separation was achieved by 120 min gradients. Peptides were loaded and eluted with a nonlinear gradient of increasing buffer B (0.1% formic acid and 80% acetonitrile) and decreasing buffer A (0.1% formic acid) at a flow rate of 300 nL/min. Buffer B was increased slowly from 5% to 30% over 95 min and ramped to 60% over 5 min, up to 95% over 5 min and held for another 5 min before reducing to 5% for column re-equilibration for 5 min. Sample acquisition was randomized to avoid bias. Mass spectrometric analysis was performed as described in Brunner *et al.*(39), either in data-dependent (ddaPASEF) or data-independent (diaPASEF) mode. The ddaPASEF method was used for library generation. For ddaPASEF, 1 MS1 survey TIMS-MS and 10 PASEF MS/MS scans were acquired per acquisition cycle. Ion accumulation and ramp time in the dual TIMS analyzer was set to 100 ms each and the ion mobility range was analyzed from 1/K0 = 1.6 Vs cm^-2^ to 0.6 Vs cm^-2^. Precursor ions for MS/MS analysis were isolated with a 2 Th window for m/z < 700 and 3 Th for m/z >700 in a total m/z range of 100-1.700 by synchronizing quadrupole switching events with the precursor elution profile from the TIMS device. Collision energy was lowered linearly as a function of increasing mobility starting from 59 eV at 1/K0 = 1.6 VS cm^-2^ to 20 eV at 1/K0 = 0.6 Vs cm^-2^. Singly charged precursor ions were excluded with a polygon filter (Bruker Daltonik GmbH). Precursors for MS/MS were picked at an intensity threshold of 1.000 AU and re-sequenced until reaching a ‘target value’ of 20.000 AU considering a dynamic exclusion of 40 s elution. For diaPASEF analysis, the correlation of Ion Mobility (IM) with m/z was used and the elution of precursors from each IM scan was synchronized with the quadrupole isolation window. The collision energy was ramped linearly as a function of the IM from 59 eV at 1/K0 = 1.6 Vs cm^−2^ to 20 eV at 1/K0 = 0.6 Vs cm^−2^.

#### Spectronaut Data Processing

Data analysis of mass spectrometric raw files acquired in ddaPASEF mode were analyzed with Spectronaut Pulsar (version 2.4, Biognosys AG, Schlieren, Switzerland) to generate the library using the default settings. The library consisted of 175,301 precursors, 115,684 peptides, 10,191 protein groups. The ddaPASEF library were used in the targeted analysis of diaPASEF data for the CNS data against the Mus musculus UniProt fasta database (2019 release, UP00000589_10090). Peptide spectral match (PSM) and protein level FDR of 1%. A minimum of seven amino acids was required including N-terminal acetylation and methionine oxidation as variable modifications and carbamidomethylation as a fixed modification. Enzyme specificity was set to trypsin/P with a maximum of two allowed missed cleavages. Peptide identifications by MS/MS were transferred by matching four-dimensional isotope patterns between the runs (MBR) with a 0.7 min retention time match window and a 0.05 1/K0 ion mobility window. diaPASEF analysis was conducted with the Spectronaut software (v14.8, Biognosys AG, Schlieren, Switzerland) under default settings. Search parameters were according to default settings. ‘Cross run normalization’ was enabled with strategy of ‘local normalization’ based on rows with ‘Qvalue complete’. FDR was set to 1% at peptide precursor level and 1% at protein level. Decoy hits and proteins, which did not pass the Q-value threshold, were filtered out prior to data analysis.

#### MS Data Analysis and Visualization

Quantitative protein abundance data generated in Spectronaut was analyzed in a Python-based version of the Clinical Knowledge Graph (CKG) (v1.0b1 BETA, https://CKG.readthedocs.io)(40), where data handling including filtering, normalization, annotation, statistics, enrichment analysis, and visualization were conducted. A minimum of 3 quantified values out 4 replicates in at least one group were required when analyzing the data. Missing values were imputed using a mixed model of k-Nearest Neighbor’s imputation method (KNN) which assumes that the values are Missing Completely At Random (MCAR), and Probabilistic Minimum Imputation approach (MinProb) for missing values that are considered Missing Not At Random (MNAR) (downshift of 2.4 standard deviations and a width of 0.3 standard deviations). Protein intensities were log2-transformed for further analysis. For the two-samples *t*-tests, each region was analyzed separately to eliminate noise from imputation and to achieve high stringency and confidence. For generation of the abundance distribution curves, median protein abundances across all samples within a proteome were used. Coefficients of variation (CVs) were calculated and reported as overall coefficient of variation. Correlations between LFQ intensities within biological replicates and regions were done by Pearson correlations. The subsequent data analysis includes a dimensionality reduction step to enable visualization of the high dimensional proteomic datasets using two or three-dimensional representations. Linear dimensionality reduction (Principal Component Analysis (PCA)) was implemented. Statistical tests across 4 groups (2 genotypes and 2 regions) were performed by a one-way ANOVA using Benjamini-Hochberg False Discovery Rate (BH-FDR) to correct for multiple hypothesis testing. All *t*-tests performed were two-sided, unpaired, and corrected for multiple testing. Volcano plots were generated using FDR < 0.05.

### Functional metabolic mapping

Metabolic mapping using isotope tracing of ^13^C enriched substrates of acutely isolated brain slices were performed as previously described (41–43). Briefly, male Wt or *Ifnar1^−/−^* mice were euthanized by cervical dislocation after which the brain was quickly removed and microdissected in ice-cold artificial cerebrospinal fluid (ACSF) into cortical and hippocampal sections. The dissected sections were sliced (350 µm) on a McIlwain tissue chopper (The Vibratome Company, O’Fallon, MO, USA). The slices were submersed in ACSF containing either [U-^13^C]glucose (5 mM), [1,2-^13^C]acetate (5 mM), [U-^13^C]glutamate (0.2 mM), or [U-^13^C]glutamine (0.2 mM) and incubated for 1 h. The ^13^C enrichment of metabolites in tissue extracts were determined by gas chromatography-mass spectrometry (GC-MS). Data is presented as the cycling ratio, describing the rate of ^13^C accumulation, i.e. reflecting the rate of the TCA cycle (41, 44). High-performance liquid chromatography (HPLC) was used to assess amino acid amounts in slices and was performed as previously described (41, 44).

### Behavior

All behavior tests were performed and analyzed with the experimenter blinded to genotype. Mice were habituated to the behavior room 24 hours prior to testing. Behavior of *Ifnar1^−/−^* mice were evaluated against Wt and *Ifnb^−/−^* mice. Behavior of Syn1^Cre^;*Ifnar1^fl/fl^*mice were evaluated against Syn1^Cre^;*Ifnar1^+/fl^* and Syn1^Cre^;*Ifnar1^+/+^* littermates. Behavior of GFAP^Cre^;*Ifnar1^fl/fl^*mice were evaluated against GFAP^Cre^;*Ifnar1^+/fl^* and GFAP^Cre^;*Ifnar1^+/+^* littermates. Ethovision XT v11 software (Noldus) was used to measure performances on the open field (OF), elevated plus maze (EPM), Morris water maze (MWM), and Barnes maze. To minimize potential confounding stress of conducting multiple behavior tests during one time point, anxiety-like behavior assessments (OF and EPM) were conducted first, followed by motor (wire suspension and rotarod) tests, cognitive measures (Barnes maze and MWM), and pain sensitivity (tail-flick) tests (12). The novelty-suppressed feeding test and sucrose preference test assessing depression-like phenotypes were conducted on separate cohorts.

Wire suspension, rotarod, MWM, and tail-flick tests were performed as previously described (13). EPM and Barnes maze were performed as described in (12). OF test was performed as described in (45).

Novelty-suppressed feeding test was adapted from (46) and (47). Mice were fasted 24 h prior to the test, then assessed individually for 5 min on their behavior when given sudden access to a food pellet of known weight in a novel open arena. Reduced time spent eating and food intake were measured as reduced motivation and depression-like behaviors. Six mice (two Wt and four *Ifnar1^−/−^*) did not eat throughout the test duration and were thus excluded from analysis.

Sucrose preference test was adapted from (48). Mice were given free access to both water and a solution containing 1.5% weight/volume sucrose for seven consecutive days. Total liquid intake, sucrose preference ratio, and total sucrose intake were measured at the same timepoint each day. Depression-like behavior was measured as a reduced preference a rewarding activity (i.e. sucrose consumption).

Rearing/climbing test was adapted from (49). Mice were placed in a 10 x 15 cm-tall wire mesh cylinder, and video was recorded for 5 min. Rearing activity was defined as both forepaws stretched above the midline of the mouse and on the apparatus. Climbing activity was defined as all four paws lifted off the ground. Number of rearing and climbing events and cumulative time spent rearing or climbing were assessed as measures of spontaneous motor activity.

### Microscope imaging and immunoblotting

Immunofluorescence (IF), Immunohistochemistry (IHC), Transmission Electron Microscopy (TEM), and immunoblotting were performed using previously described methods (12, 13, 50). Primary antibodies used in this study are listed in Table 2.

**Table 2.**
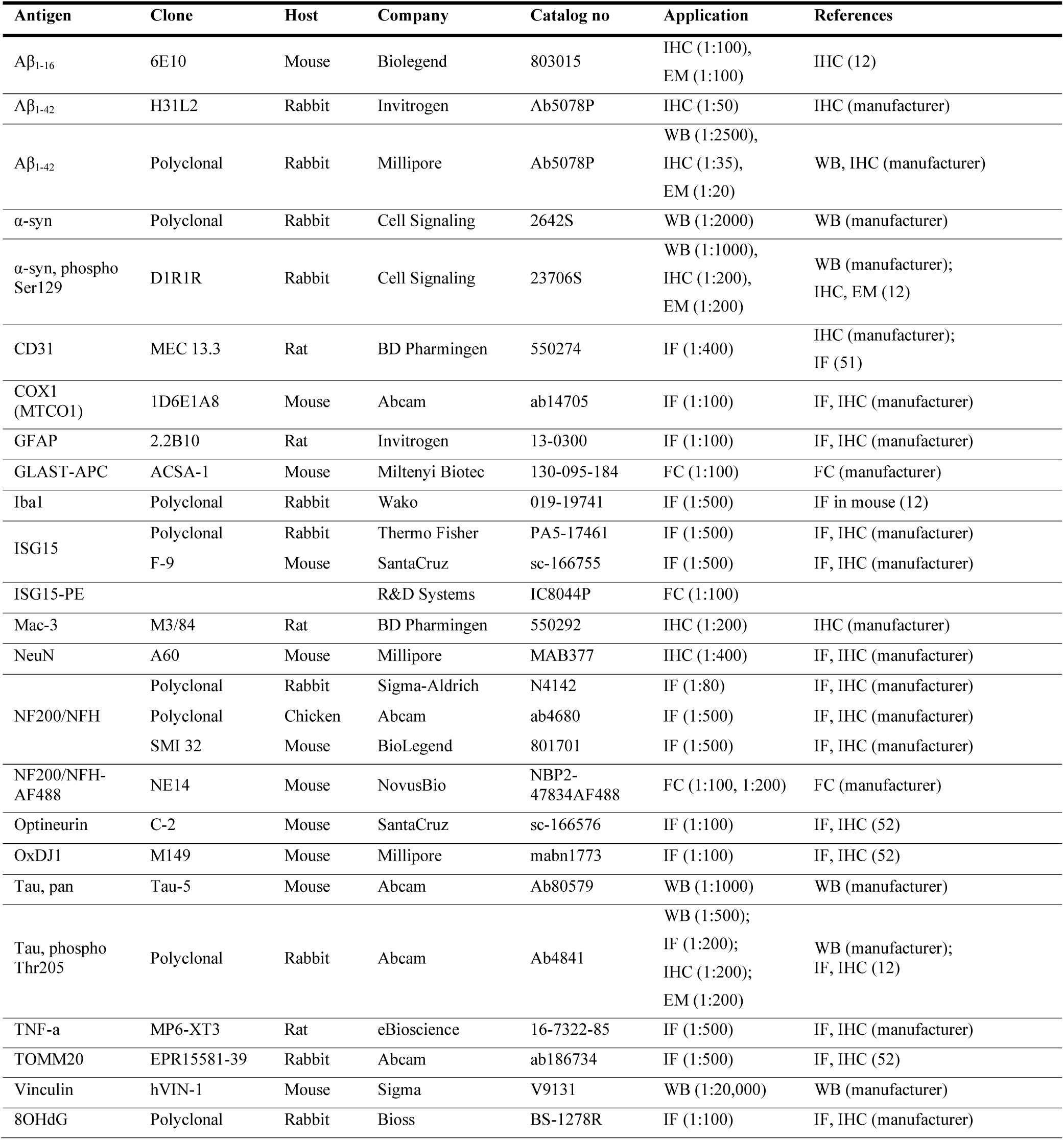
Primary antibody uses and concentrations in this study.

| Antigen | Clone | Host | Company | Catalog no | Application | References |
| --- | --- | --- | --- | --- | --- | --- |
| Aβ<sub>1-16</sub> | 6E10 | Mouse | Biologend | 803015 | IHC (1:100),<br>EM (1:100) | IHC (12) |
| Aβ<sub>1-42</sub> | H31L2 | Rabbit | Invitrogen | Ab5078P | IHC (1:50) | IHC (manufacturer) |
| Aβ<sub>1-42</sub> | Polyclonal | Rabbit | Millipore | Ab5078P | WB (1:2500),<br>IHC (1:35),<br>EM (1:20) | WB, IHC (manufacturer) |
| α-syn | Polyclonal | Rabbit | Cell Signaling | 2642S | WB (1:2000) | WB (manufacturer) |
| α-syn, phospho<br>Ser129 | D1R1R | Rabbit | Cell Signaling | 23706S | WB (1:1000),<br>IHC (1:200),<br>EM (1:200) | WB (manufacturer);<br>IHC, EM (12) |
| CD31 | MEC 13.3 | Rat | BD Pharmingen | 550274 | IF (1:400) | IHC (manufacturer);<br>IF (51) |
| COX1<br>(MTCO1) | 1D6E1A8 | Mouse | Abcam | ab14705 | IF (1:100) | IF, IHC (manufacturer) |
| GFAP | 2.2B10 | Rat | Invitrogen | 13-0300 | IF (1:100) | IF, IHC (manufacturer) |
| GLAST-APC | ACSA-1 | Mouse | Miltenyi Biotec | 130-095-184 | FC (1:100) | FC (manufacturer) |
| Iba1 | Polyclonal | Rabbit | Wako | 019-19741 | IF (1:500) | IF in mouse (12) |
| ISG15 | Polyclonal | Rabbit | Thermo Fisher | PA5-17461 | IF (1:500) | IF, IHC (manufacturer) |
|  | F-9 | Mouse | SantaCruz | sc-166755 | IF (1:500) | IF, IHC (manufacturer) |
| ISG15-PE |  |  | R&D Systems | IC8044P | FC (1:100) |  |
| Mac-3 | M3/84 | Rat | BD Pharmingen | 550292 | IHC (1:200) | IHC (manufacturer) |
| NeuN | A60 | Mouse | Millipore | MAB377 | IHC (1:400) | IF, IHC (manufacturer) |
| NF200/NFH | Polyclonal | Rabbit | Sigma-Aldrich | N4142 | IF (1:80) | IF, IHC (manufacturer) |
|  | Polyclonal | Chicken | Abcam | ab4680 | IF (1:500) | IF, IHC (manufacturer) |
|  | SMI 32 | Mouse | BioLegend | 801701 | IF (1:500) | IF, IHC (manufacturer) |
| NF200/NFH-AF488 | NE14 | Mouse | NovusBio | NBP2-47834AF488 | FC (1:100, 1:200) | FC (manufacturer) |
| Optineurin | C-2 | Mouse | SantaCruz | sc-166576 | IF (1:100) | IF, IHC (52) |
| OxDJ1 | M149 | Mouse | Millipore | mabn1773 | IF (1:100) | IF, IHC (52) |
| Tau, pan | Tau-5 | Mouse | Abcam | Ab80579 | WB (1:1000) | WB (manufacturer) |
| Tau, phospho<br>Thr205 | Polyclonal | Rabbit | Abcam | Ab4841 | WB (1:500);<br>IF (1:200);<br>IHC (1:200);<br>EM (1:200) | WB (manufacturer);<br>IF, IHC (12) |
| TNF-α | MP6-XT3 | Rat | eBioscience | 16-7322-85 | IF (1:500) | IF, IHC (manufacturer) |
| TOMM20 | EPR15581-39 | Rabbit | Abcam | ab186734 | IF (1:500) | IF, IHC (52) |
| Vinculin | hVIN-1 | Mouse | Sigma | V9131 | WB (1:20,000) | WB (manufacturer) |
| 8OHdG | Polyclonal | Rabbit | Bioss | BS-1278R | IF (1:100) | IF, IHC (manufacturer) |

### In situ hybridization assay

The *in-situ* hybridization assay RNAscope (ACDbio) was used to detect neuronal and astrocytic *Ifnar1* expression in mouse brain, which was performed according to the manufacturers protocol with the Mm-*Ifnar1* probe (cat# 512971-C2) and RNAscope Multiplex Fluorescent Reagent Kit v2 (cat# 323100). In brief, paraffin embedded coronal brain slices were deparaffinized with Xylene and ethanol washes (99%), followed by hydrogen peroxide treatment. Antigen retrieval was conducted with the supplied solution by microwave boiling, followed by protease treatment (supplied with the kit). The hybridization with mm-*Ifnar1* probe was done in the HyBEZ oven at 40C for 15 min, followed by 3 amplification steps (AMP1, AMP2 and AMP3) at 40C 30 mins each, and treatment with HRP-C1for 15min at 40C. Finally, slides were incubated with the fluorescent label for the probe (Opal570) for 30min at 40C, followed by HRP blocker for 15mins at 40°C. Conventional immunofluorescent staining with antibodies against cell specific markers GFAP and NF200 (outlined in Table 2) was then performed as previously described.

### Flow Cytometry (FC)

All kits were used according to manufacturer’s protocols. Briefly, snap-frozen brains from 3-month-old Wt*, Ifnb^−/−^*and *Ifnar1^−/−^* mice (n=4, 2M and 2F per genotype) were prepared in a single cell suspension using the Invent Biotechnologies inc. Minute™ Cell Suspension Isolation Kit from Fresh/Frozen Tissues (#CS-031). For myelin removal, a 33% Percoll gradient was run at 800 *g* for 15 min (low break). The single cells suspension was counted and stained using the antibodies outlined in Table 2. Briefly, cells were first stained with LIVE/DEAD™ Fixable Violet Dead Cell Stain Kit, for 405 nm excitation (#L34955) and blocked with an BD Fc block (#553142, 1:200). Next, cells were stained for surface markers (GLAST) for 30 min at 4°C, after which cells were fixed and permeabilized using BD Cytofix/Cytoperm™ Fixation/Permeabilization Kit (#554714) and stained for intracellular epitopes (NF200 and ISG15) for 30 min at 4°C. Compensation using UltraComp eBeads™ Compensation Beads (01-2222-41) and ArC™ Amine Reactive Compensation Bead Kit (# A10346). The samples were analyzed by the BD LSRFortessa™ X-20 Cell Analyzer with HTS. The analysis was performed in FlowJo 10.8.2, using fluorescence minus one (FMO controls) for gating.

### Quantitative PCR (qPCR)

All kits were used according to manufacturer’s protocols. For validation of *Ifnar1* knockdown, primary neurons and astrocytes were isolated from cortices of P0/P1 Syn1^Cre^;*Ifnar1^fl/fl^*, GFAP^Cre^;*Ifnar1^fl/fl^*, and control Cre negative *Ifnar1fl/fl* pups (n=3-7 per genotype) and cultured as previously described (13, 50). For investigation of immune cytokines, snap-frozen cortices from 3-month-old Wt*, Ifnb^−/−^* and *Ifnar1^−/−^* mice (n=3, 2M and 1F per genotype) were used. Briefly, RNA was isolated from snap-frozen cortices of or primary neurons and astrocyte cultures using Qiagen AllPrep kit (#80004). Biorad iScript™ cDNA Synthesis Kit (#1708891) was used for reverse transcription of RNA in complementary DNA (cDNA), using 350 ng of template RNA. For the qPCR reaction Roche Faststart SYBR green master (#4673484001) was used on 10 ng of cDNA and 10 µM primers (Table 3) per reaction. Technical replicates averages were used for subsequent analysis, standardizing results to two housekeeping genes (*Beta Actin* and *Rpl13a*). Data were further normalized to the average of Wt replicates, shown as fold change. Statistical tests used are outlined in the figure legend of each experiment.

**Table 3:**
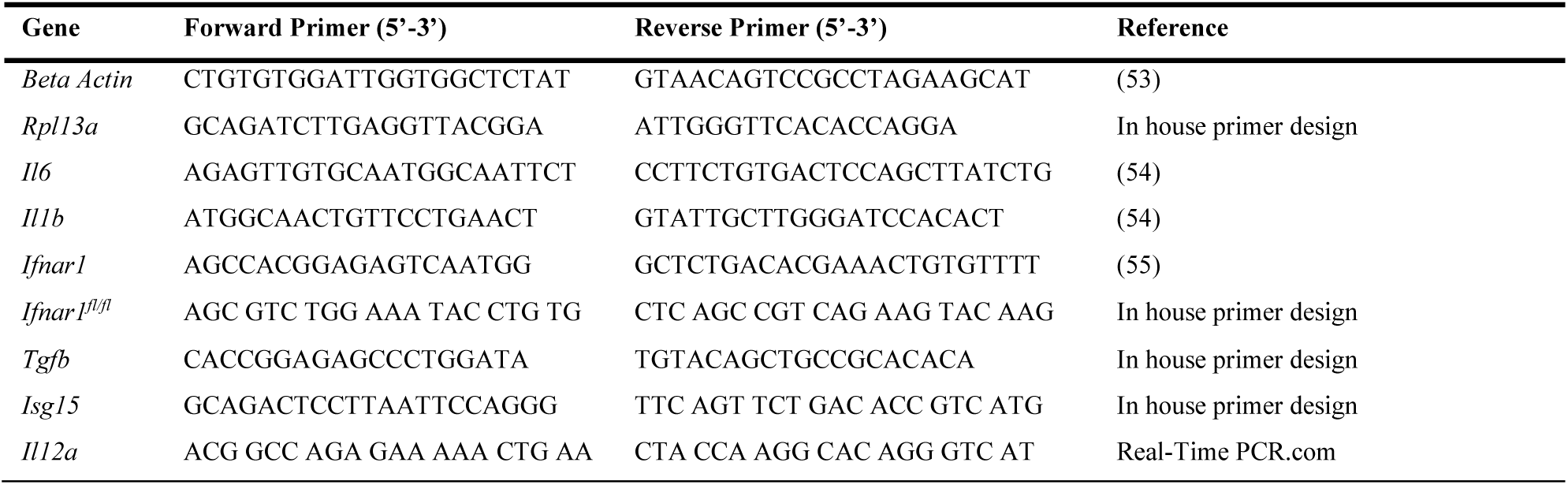
qPCR Primers used in the study.

### Statistical Analysis

Prism software (v9) was used for statistical analysis of results from behavior tests, FC, qPCR, immunohistochemistry, immunoblotting, and GC-MS metabolic labelling experiments. Parametric tests were applied based on passing the Shapiro-Wilk normality test. Unpaired Student’s *t*-tests were used for two-group comparisons, and one- or two-way ANOVAs were applied to multiple-group comparisons. Statistical tests used for each experiment are specified in the figure legends. *P* values <0.05 were considered statistically significant.

## Results

### *IFNAR1* is differentially expressed in human brain cell types and is affected in patients with PD and LBD

Though we previously reported that Type I-IFN signaling is dysfunctional in PD patients upon dementia progression (10), the primary cellular sources of general *IFNAR1* expression in the entire brain had not been identified; thus, baseline cellular expression patterns of *IFNAR1* in the human brain were examined. 10x snRNA-seq expression data from the Allen Brain Institute (15) revealed differential *IFNAR1* expression among the major human brain cell types, including excitatory and inhibitory neurons, astrocytes, oligodendrocytes, oligodendrocyte precursor cells (OPCs), and microglia (Fig. 1A). In general, astrocytic *IFNAR1* expression was notably lower compared to neurons and the other brain cell types. Wild-type (Wt) mice also demonstrated significantly lower astrocytic *Ifnar1* expression compared to neurons using *in situ* hybridization combined with immunofluorescent staining using GFAP vs NF200 for visualization of astrocytic vs. neuronal *Ifnar1* expression, respectively (Fig. 1B). We next used a publicly available single cell RNA-seq dataset (16) to compare PD and LBD cortical samples to unaffected controls and determined how *IFNAR1* and associated Type-I IFN signaling gene (10) expression were affected among different brain cells with or without dementia (Fig. 1C). Notably, *IFNAR1* is specifically lower in LBD vs control and PD in dopaminergic (DA) neurons and microglia, which supports our previous observations in PD and PDD cortical samples where *IFNAR1* expression was significantly reduced in PDD versus PD (10). IFNβ, lack of which we previously observed to result in PD and LBD-like progression in mice, had reduced expression in both LBD and PD dopaminergic neurons compared to controls (13). *IFNAR1* and several related genes were also reduced in astrocytes, oligodendrocytes, and OPCs in both LBD and PD compared to controls, implicating *IFNAR1* signaling dysregulation in neurodegenerative mechanisms involved in both neurodegenerative conditions.

**Fig. 1.**
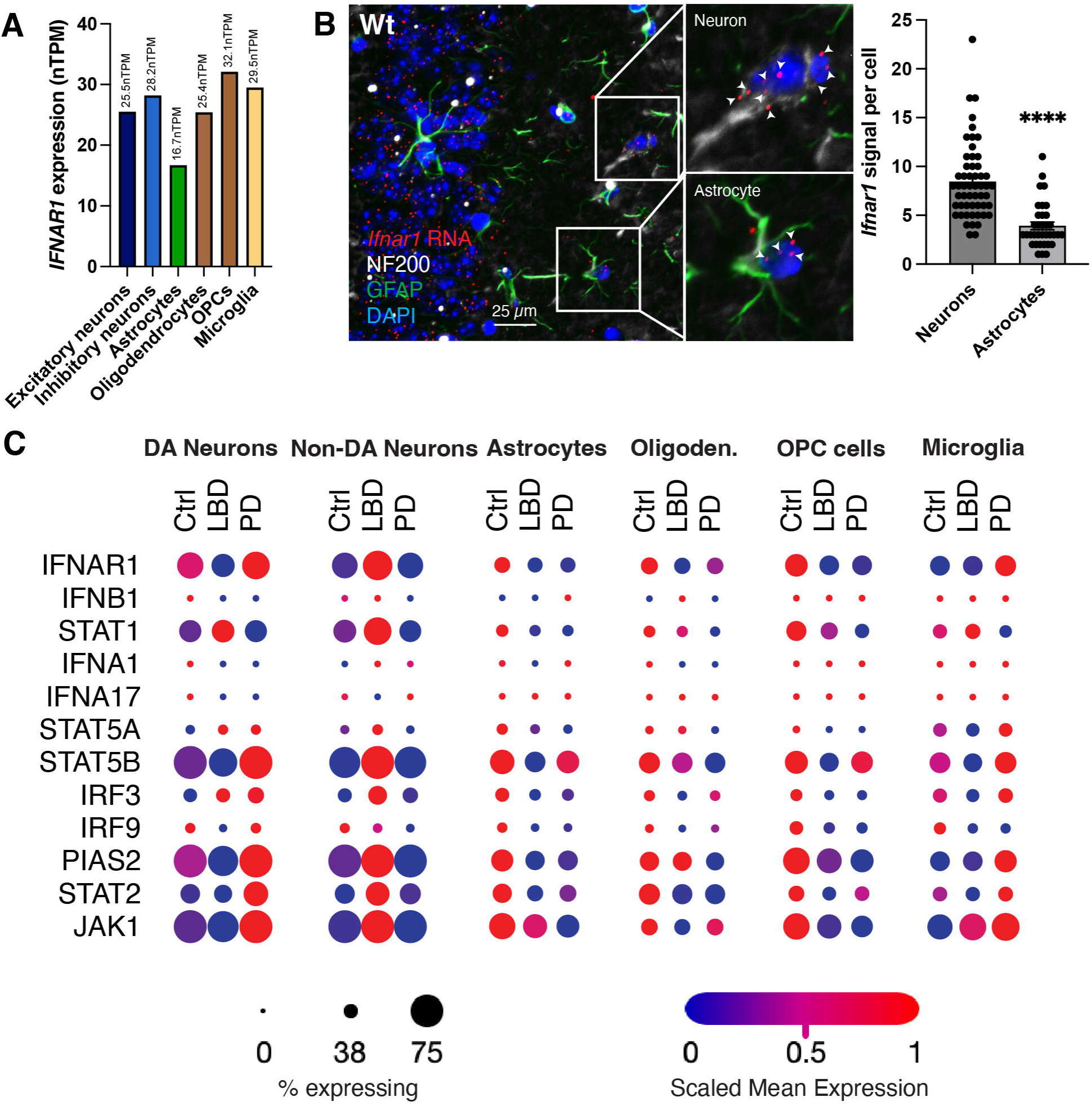
*IFNAR1* expression in human brain varies by cell type and is differentially affected in PD and LBD. (**A**) *IFNAR1* RNA expression in human brain cell types. OPCs = Oligodendrocyte Precursor Cells. (**B**) Differential RNA expression of *Ifnar1* (red) in wild-type (Wt) mouse NF200^+^ neurons (white; *n =* 54) and GFAP^+^ astrocytes (green; *n =* 35) using *in situ* hybridization. DAPI = blue. \*\*\*\**P* < 0.0001 by unpaired *t*-test. Scale bar, 25 µm. (**C**) Dot plot comparing *IFNAR1* and related Type-I IFN gene expression in different brain cells from an single cell RNA-seq dataset (16) derived from human PD and LBD patients compared to unaffected controls (*n*=3-8). DA = Dopaminergic. Scaled mean expression = relative gene expression across the annotated cell type within each patient category.

These results show that *IFNAR1* expression is subject to varying regulation across different brain cell types in both healthy human and mouse brains, warranting further investigation into the relationship between brain cell-specific changes in *IFNAR1* expression and dementia development.

### Single nuclei (sn)RNA-seq in *Ifnar1^−/−^* brain revealed PD-related changes in neuronal and astrocytic mitochondrial genes and neurotransmission

To identify potential brain cell-specific changes upon IFNAR1 loss *in vivo*, snRNA-seq was utilized on cortical nuclei isolated from young adult (1.5-month-old) Wt and *Ifnar1^−/−^*mice. Cortical cell types were grouped into five broad cell classes based on their gene expression profile (Fig. 2A; Supplementary Fig. 1A). Glutamatergic neurons (Glut) had the highest number of significant (*P <* 0.05) differentially expressed (DE) genes (396), followed by oligodendrocytes (Oligo, 294), GABAergic neurons (GABA, 242), microglia (Micro, 206), and astrocytes (Astro, 52) (Fig. 2B). Unbiased hierarchical clustering of gene expression showed tight clustering of biological replicates within genotypes for each cell class (Supplementary Fig. 1B).

**Fig. 2.**
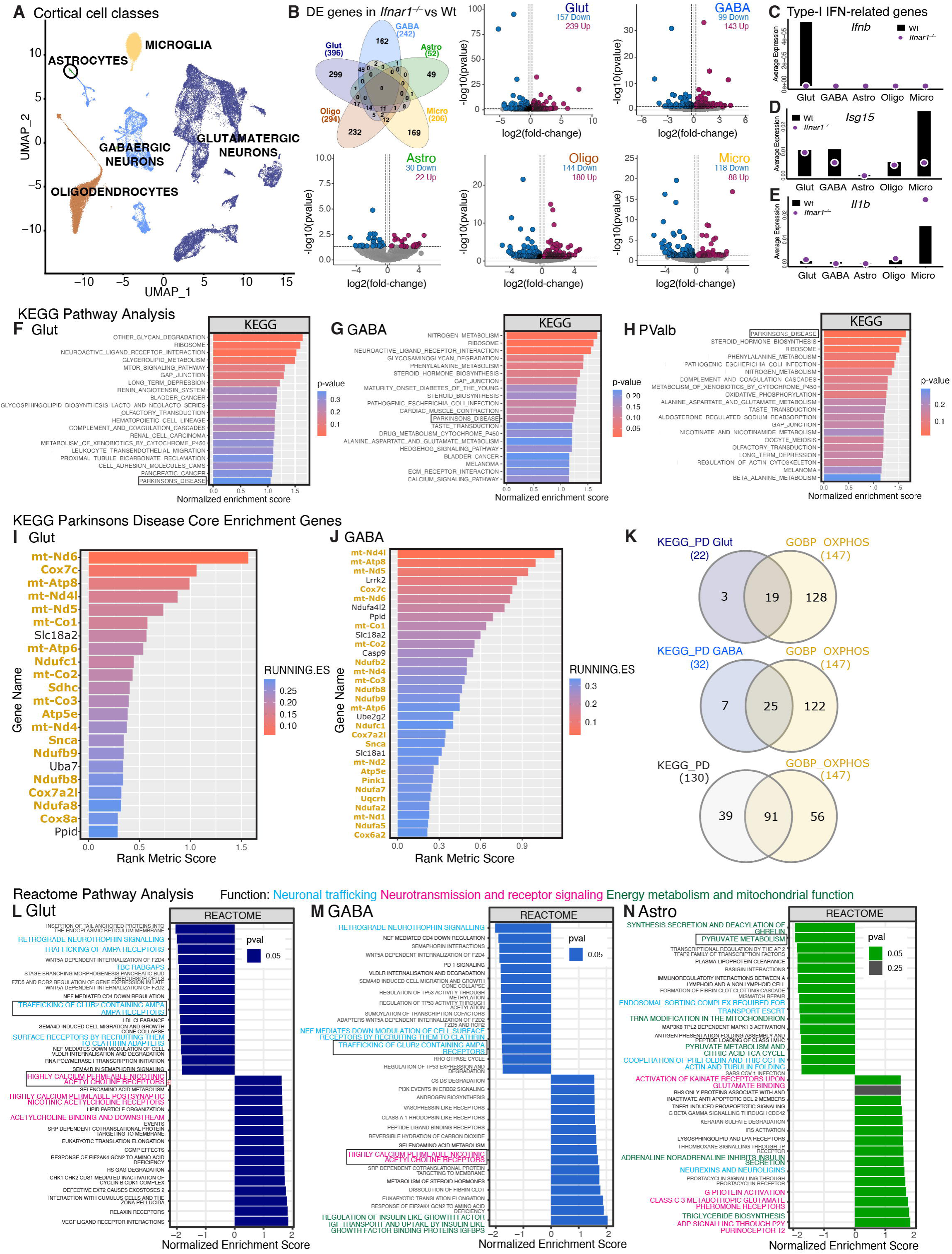
snRNA-seq revealed early PD-related changes in neuronal mitochondrial genes, neurotransmission, and glial support upon *Ifnar1^−/−^*. (**A**) Uniform manifold approximation and projection (UMAP) of broad cell classes isolated from cortex from 1.5-month-old Wt vs *Ifnar1^−/−^* mice, *n* = 4 per genotype. A total of 26020, 22821, 24268, 21069, and 15185 genes identified in glutamatergic neurons, GABAergic neurons, astrocytes, microglia, and oligodendrocytes, respectively. (**B**) Venn diagram showing overlap of significant (P < 0.05) differentially expressed (DE) genes identified in glutamatergic neurons (Glut), GABAergic neurons (GABA), astrocytes (Astro), oligodendrocytes (Oligo), and microglia (Micro), and volcano plots for each cell class showing significance and fold-change (FC) of DE genes. (**C-E**) Specific expression of interferon-related genes (**C**) *Ifnb,* (**D**) *Isg15*, and (**E**) *Il1b* in each cell class. Purple circles represent *Ifnar1^−/−^* and black bars represent Wt average expression. (**F-H**) Top 20 upregulated KEGG pathways for (**F**) glutamatergic neurons, (**G**) GABAergic neurons, and (**H**) parvalbumin inhibitory neurons, with KEGG ‘Parkinsons Disease’ pathway indicated in a black box. (**I-J**) Core enrichment genes within the KEGG ‘Parkinsons Disease’ (KEGG_PD) pathway for (**I**) glutamatergic neurons and (**J**) GABAergic neurons, with mitochondrial genes (belonging to the ‘GOBP Oxidative Phosphorylation’ (GOBP_OXPHOS) pathway) labelled in yellow. (**K**) Overlap between KEGG_PD core enrichment genes for glutamatergic neurons, GABAergic neurons, and the KEGG_PD pathway overall with GOBP_OXPHOS genes. (**L-M**) GSEA-Reactome enrichment tables showing top 15 positively and negatively enriched pathways for (**L**) glutamatergic neurons, (**M**) GABAergic neurons, and (**N**) astrocytes. Pathways relevant to neuronal and astrocytic function are highlighted, where blue = trafficking affecting neurotransmission; magenta = neurotransmission and receptor signaling; green = energy metabolism and mitochondrial function. Pathways enclosed in a black box have heatmaps featured by cell type in Supplementary Fig. 2D-G.

Next, gene set enrichment analysis (GSEA) was conducted on all cell classes to examine genotypic differences. Expression of interferon-related genes in each cell class was assessed to determine the validity of the experimental approach. Glutamatergic and GABAergic neuronal classes expressed dysregulated type-I IFN or related pathways involving inflammation and cytokine receptor signaling, inclusive of IFN-α, IFN-γ, IL-6, TNF-α, and TGF-β (Supplementary Fig. 1C). Glutamatergic neurons had the highest expression of *Ifnb* in Wt mice, which was reduced in *Ifnar1^−/−^* compared to Wt (Fig. 2C)*. Isg15* (interferon-stimulated gene 15), an E3 ubiquitin ligase directly regulated by IFNAR1 signaling (56), was also downregulated in general in *Ifnar1^−/−^* cortical cells (Fig. 2D), which was confirmed with qPCR and FC in *Ifnar1^−/−^*brains (Supplementary Fig. 1D, E). Proinflammatory interleurkin-1beta (*Il-1b*), normally is downregulated by IFNAR1 signaling (57), was upregulated in *Ifnar1^−/−^* microglia (Fig. 2E), suggesting potential neuroinflammation and has been associated with dysregulated Type-I IFN signaling PDD (10). Accordingly, *Ifnar1^−/−^*microglia showed negative enrichment of interferon signaling-related Reactome and KEGG (Kyoto encyclopedia of genes and genomes) pathways, but significant positive enrichment in the KEGG pathways ‘Parkinson’s disease’, ‘Alzheimer’s disease’, and ‘Huntington’s disease’, collectively demonstrating that *Ifnar1^−/−^*microglia adopt a neurodegenerative phenotype (Supplementary Fig. 1F).

GSEA of glutamatergic and GABAergic neuronal classes featured the KEGG pathway ‘Parkinson’s disease’ among the top 20 upregulated KEGG pathways (Fig. 2F, G), which was reflected in the majority of annotated cortical neuronal subtypes within each class (Supplementary Fig. 1B and Supplementary Fig. 2A, B), with Parvalbumin^+^ (PValb) interneurons shown as representative subtype (Fig. 2H). Notably, mitochondrial genes (belonging to the GOBP-OXPHOS pathway) were overrepresented among KEGG ‘Parkinson’s disease’ for glutamatergic (Fig. 2I) and GABAergic (Fig. 2J) neurons, making up 86% and 78% of core enrichment genes respectively, compared to 70% overall genes shared between KEGG ‘Parkinson’s disease’ and GOBP-OXPHOS (Fig. 2K). Accordingly, several mitochondrial and metabolic pathways were among the top 15 dysregulated Reactome pathways in both glutamatergic and GABAergic neuronal classes (Fig 2L, M, Supplementary Fig. 2C).

Reactome pathways related to neuronal trafficking were among the top 15 negatively enriched pathways in neurons, while neurotransmission and receptor signaling-related pathways were among the top 15 positively enriched pathways (Fig. 2L, M). Dysregulation of genes in processes relating to glutamatergic signaling and cholinergic neurotransmission (Fig. 2L, Supplementary Fig. 2D. E) suggests dysfunction of excitatory neurotransmission upon IFNAR1 loss (58). Of note, *Ifnar1^−/−^* astrocytes and oligodendrocytes also showed positive enrichment of excitatory neurotransmission pathways (Fig. 2N; Supplementary Fig. 1G); however, *Ifnar1^−/−^*astrocytes also demonstrated negative pathway enrichment relating to energy metabolism and mitochondrial function (Fig. 2N), along with several pathways involved in neurotransmission and inflammation (Supplementary Fig. 2F). Downregulation of pyruvate metabolism genes in *Ifnar1^−/−^* astrocytes and other brain energy-related processes may indicate altered brain metabolism and subsequent disturbances in neurotransmission events (Supplementary Fig. 2G).

These findings suggest that IFNAR1 loss results in cell-specific transcriptomic alterations that may collectively affect neuronal function and homeostasis. Many DE genes were unique between neuronal and astrocytic classes (Fig. 2B), suggesting distinct cell-specific dysfunctions upon IFNAR1 loss.

### Protein alterations in *Ifnar1^−/−^* brains suggest disrupted neurotransmission

To investigate how cell-specific transcriptional changes affect brain protein expression, proteome comparisons were made between Wt and *Ifnar1^−/−^*cortex and hippocampus. Evaluation of technical and biological quality showed a strong correlation between replicates (Pearson correlation ranging from 0.94–0.95) (Fig. 3A) and a clear separation of brain regions and genotypes by principal component analysis (PCA) (Fig. 3B). An average of 7657 protein groups were identified after data preparation and filtering (Supplementary Fig. 3A). Proteome coverage and dynamic range were comparable across cortex and hippocampus in both genotypes (Supplementary Fig. 3B), as well as coefficients of variation (CVs) (Wt cortex: 9.74%; *Ifnar1^−/−^*cortex: 10.97%; Wt hippocampus: 14.27%; and *Ifnar1^−/−^* hippocampus: 12.01%) (Supplementary Fig. 3C). In total, 1560 significant (FDR<0.05) protein differences were found (Fig. 3C), most of which reflected regional differences and were comparable to a previous report (42). Genotypic differences within each brain region were analyzed and visualized as differential expression on volcano plots (Fig. 3D, E).

**Fig. 3.**
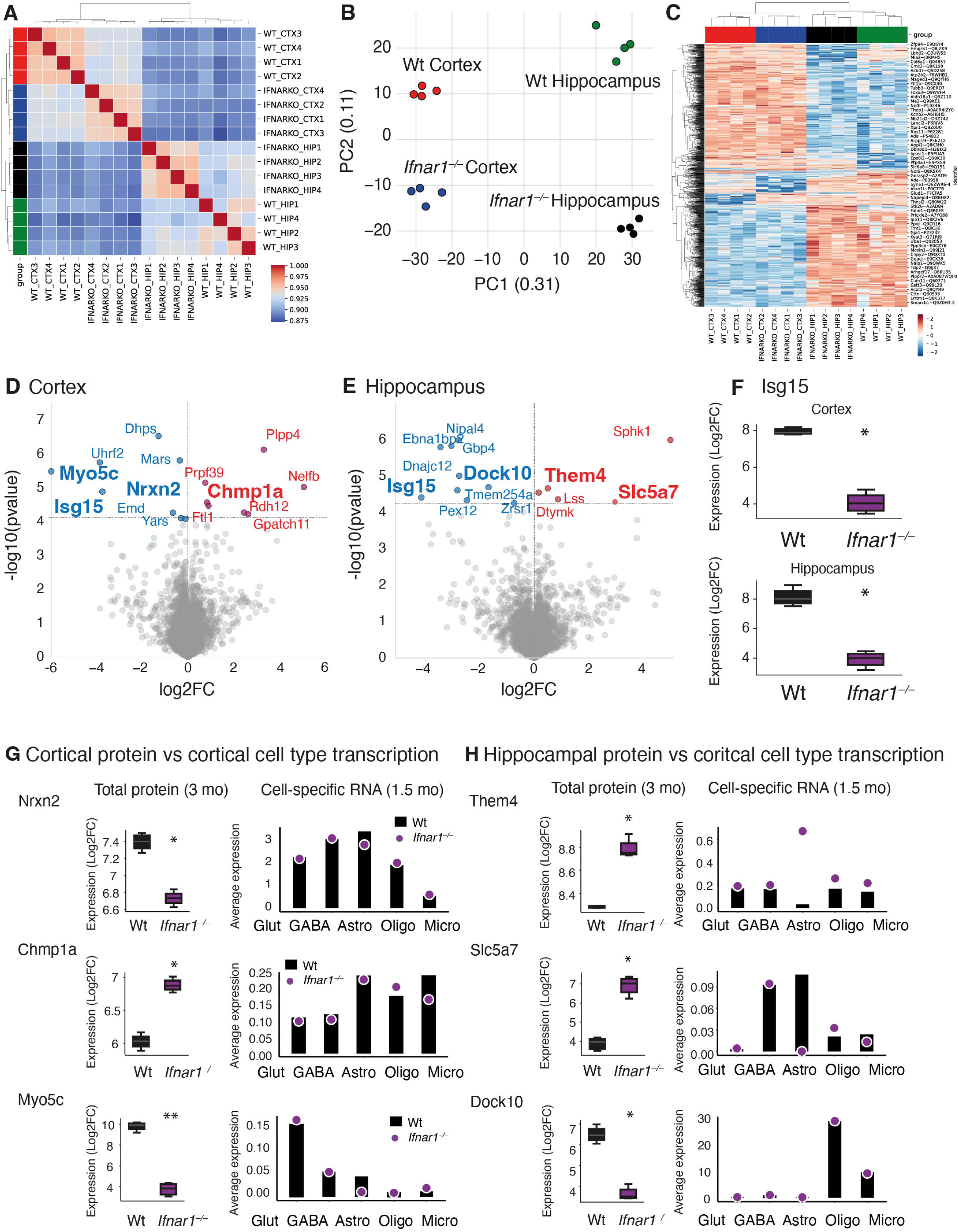
Protein alterations in *Ifnar1^−/−^* brains support snRNA-seq results suggesting dysregulated neurotransmission. (**A**) Pearson correlation plot indicating high reproducibility between samples. (**B**) Principal component analysis (PCA) showing high correlation of samples within genotype and brain region (*n* = 4 per genotype and per region). (**C**) Unbiased hierarchical clustering of differentially expressed proteins indicates distinct signatures for each brain region and some changes between genotypes within each region. (**D-E**) Volcano plots showing significant hits by false discovery rate (FDR<0.05)-adjusted p values in *Ifnar1^−/−^*vs Wt (**D**) cortex and (**E**) hippocampus. (**F**) Relative expression (Log2 FC intensity) of Isg15 in both cortex and hippocampus. (**G**) Relative protein expression of Nrxn2, Chmp1a, and Myo5c in 3-month-old *Ifnar1^−/−^*cortex compared to 1.5-month-old *Ifnar1^−/−^* cortical cell snRNA-seq expression. (**H**) Relative protein expression of Them4, Slc5a7, and Dock10 in 3-month-old *Ifnar1^−/−^* hippocampus compared to 1.5-month-old *Ifnar1^−/−^* cortical cell snRNA-seq expression. \**P* < 0.05 and \*\**P* < 0.01 by FDR-adjusted ANOVA.

Consistent with snRNA-seq, qPCR, and FC data (Fig. 2D, Supplementary Fig. 1D, E), ISG15 was significantly downregulated in both *Ifnar1^−/−^*cortex and hippocampus (Fig. 3F). Cortical changes included proteins involved in synaptic remodeling (Nrxn2, a pre-synaptic protein, reduction of which reduces N-methyl-D-aspartate (NMDA) receptor activity and spontaneous cortical neurotransmitter release (59)), endosomal sorting (Chmp1a, an essential subunit of the endosomal sorting complex required for transport (ESCRT)-III, dysfunction of which is associated with increased PDD risk (60) and pTau NFT development (61)), and microtubule dynamics (Myo5c, which facilitates actin-dependent organelle trafficking, dysregulation of which is associated with NFT manifestation (62)) (Fig. 3G). As with cortex, proteins significantly altered in *Ifnar1^−/−^* hippocampus are involved in neuronal survival, function, and synaptic regulation, including mitochondrial regulation (Them4, regulator of mitochondrial fission, mitochondrial membrane permeability, and fatty acid metabolism (63, 64)), cholinergic neurotransmission (Slc5a7, essential for synthesis of acetylcholine and mitochondria-derived acetyl-CoA (65)), and synapse remodeling (Dock10, required for synaptic morphogenesis of hippocampal neurons (66)) (Fig. 3H). Of note, protein expression and transcription patterns in both regions appeared to differ by cell type (Fig. 3G, H).

Here it was observed that cell-specific transcriptional alterations upon IFNAR1 loss affect expression of brain proteins involved in several processes required for effective neurotransmission and mitochondrial function, implicating IFNAR1 loss with general dysregulation of brain homeostasis.

### Integrated Omics revealed mitochondrial dysfunction, defective mitophagy and elevated oxidative stress upon IFNβ-IFNAR signaling loss

To identify shared and potentially central pathways disrupted upon IFNAR1 loss, analysis of cortical transcriptomic and proteomic datasets from *Ifnar1^−/−^* mice were integrated. GSEA of the proteomics dataset identified ‘Gene Ontology (GO) Cellular Component (GOCC) Mitochondrion’ and metabolic pathways as major top hits (Fig. 4A, orange and green boxes, respectively), which supported the mitochondrial and metabolic related Reactome pathways identified by GSEA in the neuronal and astrocytic transcriptomic data (Fig. 2L-N, Supplementary Fig. 2G). To directly compare cell-specific transcriptomic data with bulk proteomic data, all cellular transcriptomic data were declassified into a pseudobulk dataset. For validation of the approach, gene ontologies related to IFNAR1 depletion were checked, namely ‘GO Biological Process (GOBP) Response to Interferon beta’ (IFNb-response) and ‘GOBP regulation of viral process’ (Viral response), which appeared among the top altered GSEA pathways in the pseudobulk snRNA-seq data of *Ifnar1^−/−^* group (Fig. 4B), supporting single cell findings for related Type-I IFN genes (Fig. 2C-E) and proteomic analysis (Fig. 3F). There was also significant overlap between mitochondrial gene/protein expression and Type-1 IFN-related pathways associated with IFNβ-IFNAR responses (Fig. 4C), supporting further investigation of mitochondrial dysfunction in *Ifnar1^−/−^*mice.

**Fig. 4.**
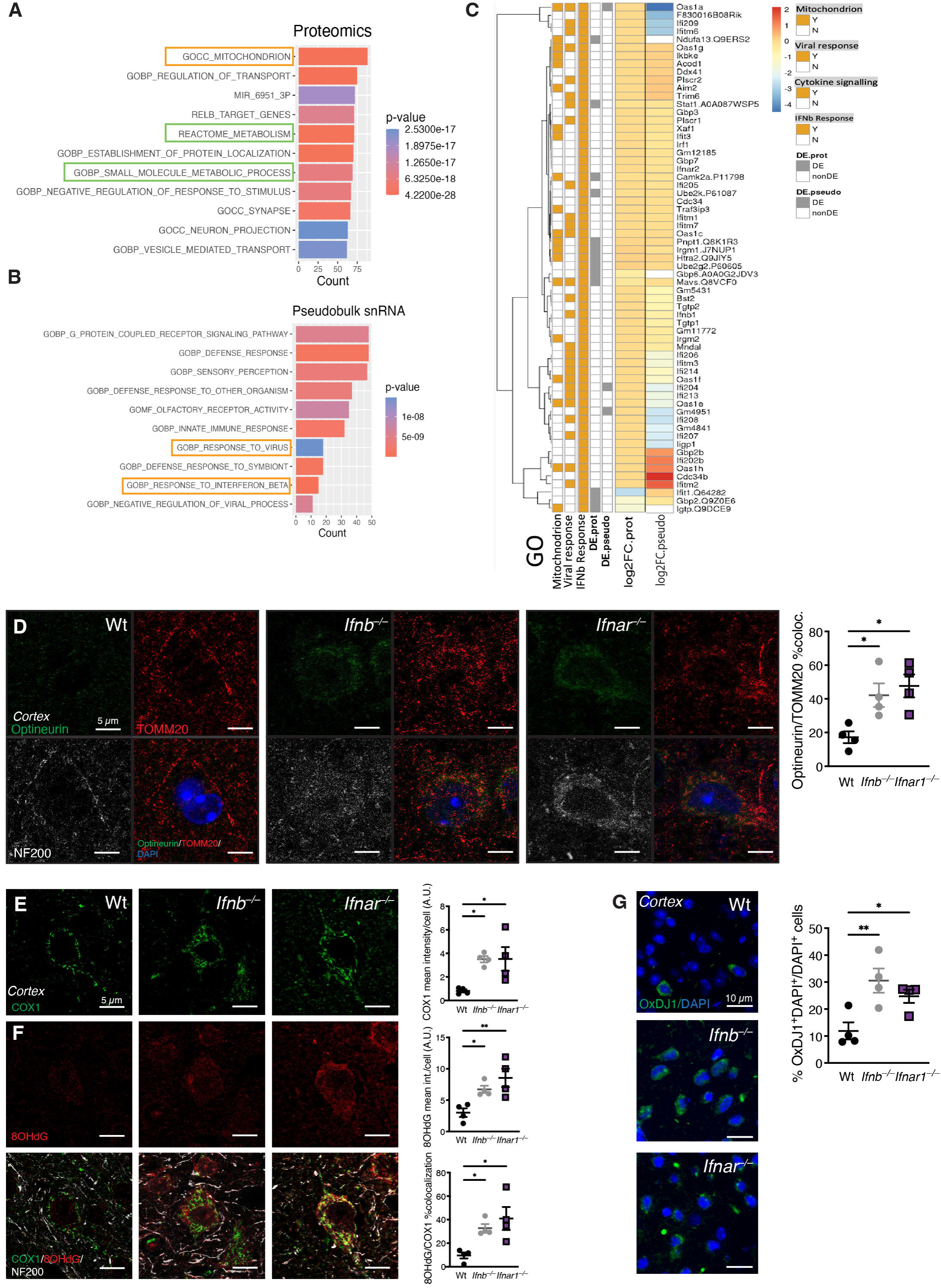
Integrated Omics revealed lack of IFNβ-IFNAR signaling results in mitochondrial dysfunction. (**A**) Top 10 GSEA pathways in the proteomics dataset with mitochondrial (yellow box) and metabolic (green boxes) pathways indicated. (**B**) Top 10 GSEA pathways in the proteomics dataset with Type-I IFN-related (yellow boxes) pathways indicated. (**C**) Heatmap indicating yes/no (Y/N) overlap and differential expression (DE) of proteins from the proteomic dataset (DE.Prot) and genes from the pseudobulk snRNA transcriptomic dataset (DE.pseudo) within ‘GOCC Mitochondrion’, ‘GOBP Response to Interferon beta’, and ‘GOBP regulation of viral process’ pathways, alongside individual protein (Log2FC.Prot) and gene (Log2FC.pseudo) expression patterns. (**D**) Representative images and quantification of average Optineurin (green) and TOMM20 (Red) colocalization in NF200^+^ (white) cortical neurons of 6-month-old Wt, *Ifnb^−/−^*, and *Ifnar1^−/−^* mice (*n* = 3-4 mice per genotype). \**P* < 0.05 by one-way ANOVA and Dunnett’s post hoc correction test. Scale bars, 5 µm. (**E-F**) Representative images and quantification of (**E**) COXI (green) mean intensity, (**F**) 8OHdG (red) mean intensity, and COXI/8OHdG %colocalization in NF200+ (white) cortical neurons of 6-month-old Wt, *Ifnb^−/−^*, and *Ifnar1^−/−^* mice (*n* = 3-4 mice per genotype). \**P* < 0.05 and \*\**P <* 0.01 by one-way ANOVA and Dunnett’s post hoc correction test. Scale bars, 5 µm. (**G**) Representative images and quantification of %OxDJ1^+^(green)/DAPI^+^(blue)/total DAPI+ cells in cortex of 6-month-old Wt, *Ifnb^−/−^*, and *Ifnar1^−/−^* mice (*n* = 3-4 mice per genotype). \**P* < 0.05 and \*\**P <* 0.01 by one-way ANOVA and Dunnett’s post hoc correction test. Scale bars, 10 µm. Data in all graphs are mean ±SEM.

To characterize potential mitochondrial defects, optineurin, a receptor for removal of damaged mitochondria (67), was investigated in *Ifnar1^−/−^* mice compared to Wt and *Ifnb^−/−^* mice, as accumulation of mitochondria in *Ifnb^−/−^* brains have been associated with defects in mitophagy (52). Optineurin colocalization with mitochondrial marker TOMM20 was elevated in brains of mice lacking IFNβ-IFNAR signaling compared to Wt (Fig. 4D). Higher levels of cytochrome C oxidase I (COXI) were also observed in both *Ifnb^−/−^* and *Ifnar1^−/−^* mice (Fig. 4E), collectively indicating defects in mitophagy (52). Oxidized DJ-1 (oxDJ-1) and 8-hydroxy-2’-deoxyguanosine (8OHdG) were measured by immunohistochemistry as indicators of oxidative stress resulting from mitochondrial dysfunction (68). Increased oxDJ1 and high cytoplasmic expression of 8OhdG colocalized to COXI were found in both *Ifnar1^−/−^* and *Ifnb^−/−^* cortex (Fig. 4F, G), showing that disrupted IFNβ-IFNAR signaling results in mitochondrial dysregulation and increased oxidative stress.

Together, these data support that both independent omics datasets identified shared signaling defects and corresponding malfunctional pathways related to a lack of IFNβ-IFNAR signaling causing mitochondrial dysfunction. These data were further verified to establish that lack of type I signaling in both *Ifnar1^−/−^* and *Ifnb^−/−^* brains results in dysregulated mitochondrial homeostasis, oxidative stress and mitophagy.

### Mitochondrial defects upon *Ifnar1^−/−^* alters neurotransmitter synthesis and glucose hypermetabolism

As neurotransmission, synaptic activity, and energy metabolism are closely linked and dependent on tricarboxylic acid (TCA) cycle activity and mitochondrial function in both neurons and astrocytes, it was next determined whether the identified mitochondrial defects affected brain energy metabolism (69). Baseline measurement of amino acids related to TCA cycle in Wt and *Ifnar1^−/−^*brains were similar in cortex or hippocampus (Fig. 5A, B), indicating that potential metabolic changes in *Ifnar1^−/−^* mice reflect functional alterations due to IFNAR1 loss, rather than genotypic differences in baseline amino acid pools.

**Fig. 5.**
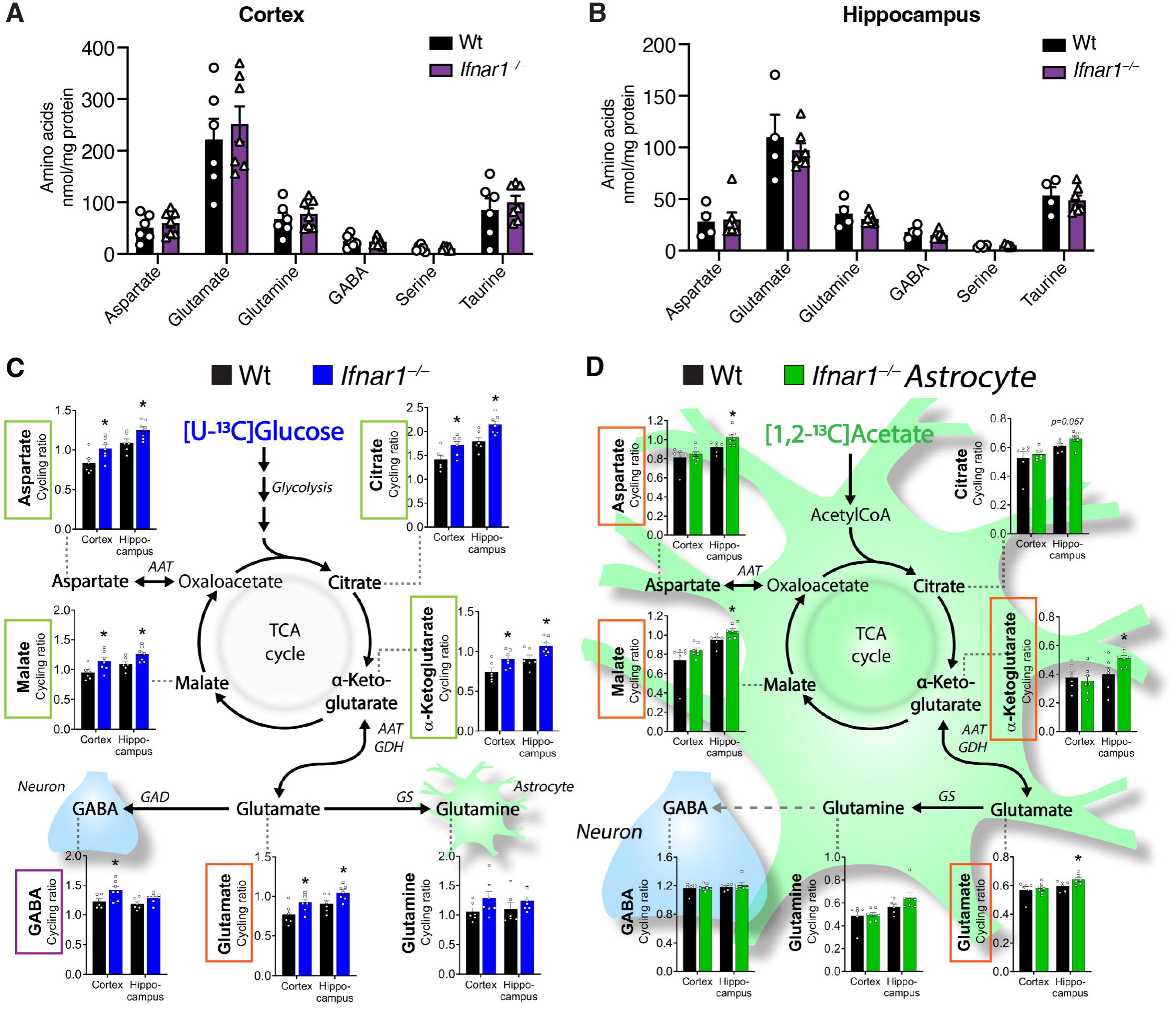
Mitochondrial defects upon *Ifnar1^−/−^* alters neurotransmitter synthesis and glucose hypermetabolism. (**A-B**) Baseline amino acid levels in (**A**) cortex and (**B**) hippocampus of Wt and *Ifnar1^−/−^*mice (*n* = 6-7 per genotype), measured by HPLC. (**C-D**) Cortical and hippocampal slices from Wt and *Ifnar1^−/−^* mice (*n* = 6-7 per genotype) were incubated with ^13^C-labelled energy substrates (**C**) [U-^13^C]glucose or (**D**) [1,2-^13^C]acetate. Cycling ratios of TCA cycle intermediates citrate, α-ketoglutarate, and malate, and amino acids aspartate, glutamine, glutamate, and GABA, in cortical and hippocampal slices, reflecting TCA cycling rate. Data are mean ±SEM. \**P* < 0.05 by *t-*test.

To functionally probe TCA cycle capacity upon IFNAR1 loss, cortical and hippocampal slices from Wt and *Ifnar1^−/−^* mice were incubated with ^13^C-labelled substrates (Fig. 5C, D; Supplementary Fig. 4A, B). Elevated metabolic activity, or hypermetabolism, in *Ifnar1^−/−^*cortex and hippocampus was revealed upon incubation with [U-^13^C]glucose, leading to significantly increased ^13^C accumulation in the TCA cycle intermediates citrate, α-ketoglutarate, malate, and the amino acid aspartate (Fig. 5C, green boxes). Increased TCA cycle activity resulted in elevated glutamate synthesis in both cortex and hippocampus (Fig. 5C, orange box), whereas GABA synthesis was significantly increased in cortex while less pronounced in hippocampus (Fig. 5C, purple box).

Though no changes were observed in [U-^13^C]glutamate or [U-^13^C]glutamine metabolism (Supplementary Fig. 4A, B), there were slight increases in ^13^C accumulation in glutamine derived from [U-^13^C]glucose metabolism (Fig. 5C). Increased ^13^C accumulation in glutamine, produced from glutamate selectively in astrocytes (41, 42), could reflect alterations in astrocyte-specific TCA cycle activity and metabolism. When probed directly with [1,2-^13^C]acetate, a substrate predominantly metabolized by astrocytes (41, 42, 70), several metabolites including aspartate, malate, α-ketoglutarate, and glutamate (Fig. 5D, orange boxes) exhibited elevated ^13^C accumulation selectively in hippocampal slices of *Ifnar1^−/−^* mice, which could reflect changes in hippocampal energy requirements from astrocytes lacking IFNAR1.

These results indicate that loss of IFNAR1 signaling results in brain glucose hypermetabolism affecting glutamate and GABA synthesis. Dysregulated excitatory neurotransmission may require greater energy demands in *Ifnar1^−/−^* neurons and astrocytes, prompting compensatory glucose metabolism and neurotransmitter production, and could indicate neuronal dysfunction and potential neurodegeneration.

### *Ifnar1***^−/−^** mice develop PDD-like neuropathology, behavior deficits, and neuroinflammation

Neuronal counts in cortex, hippocampus, and olfactory bulb were quantified to determine whether the loss of IFNAR1 signaling, and subsequent mitochondrial and metabolic dysfunction, contributes to a broad neurodegenerative phenotype similar to previous observations in *Ifnb^−/−^* brains (12, 13). Though IFNβ binds to IFNAR to conduct signaling, the lack of the *Ifnb* gene could potentially be counteracted by *Ifna* genes to compensate loss of cellular functions. Nevertheless, the number of NeuN^+^ neurons were reduced in all regions investigated (cortex, hippocampus and olfactory bulb combined) both in *Ifnar1^−/−^* and *Ifnb^−/−^* brains (Fig. 6A, Supplementary Fig. 5A-C). *Ifnar1*^−/−^ mice also exhibited neuropathological hallmarks of PDD as observed in *Ifnb^−/−^* mice (12, 13), featuring DA neuronal loss and reduction in TH^+^ neurites in the substantia nigra (Fig. 6B) with concurrent loss of TH^+^ immunoreactivity in the striatum compared to age-matched Wt mice (Fig. 6C). Notably, TH^+^ cell loss in the substantia nigra pars compacta was approximately 10% greater than loss in the ventral tegmental area (Supplementary Fig. 5D), resembling PD neuropathology (71). Significant accumulation of α-syn, phosphorylated α-syn at serine 129 (pα-syn), Tau (pan), and phosphorylated Tau at threonine 205 (pTau) was observed in aged (12-month-old) *Ifnar1^−/−^* but not Wt or young (1.5-month-old) mice (Supplementary Fig. 5E). Importantly, LB-like structures staining positive for both pα-syn and pTau were confirmed in neurons of *Ifnar1^−/−^* thalamus using both light and electron microscopy (Fig. 6D). Fibrillar structures containing Aβ were also observed by electron microscopy (Fig. 6E), and Aβ^+^ plaques like those in 5xFAD mice and *Ifnb^−/−^* mice (12) were detected in *Ifnar1^−/−^*but not Wt mice (Fig. 6F).

**Fig. 6.**
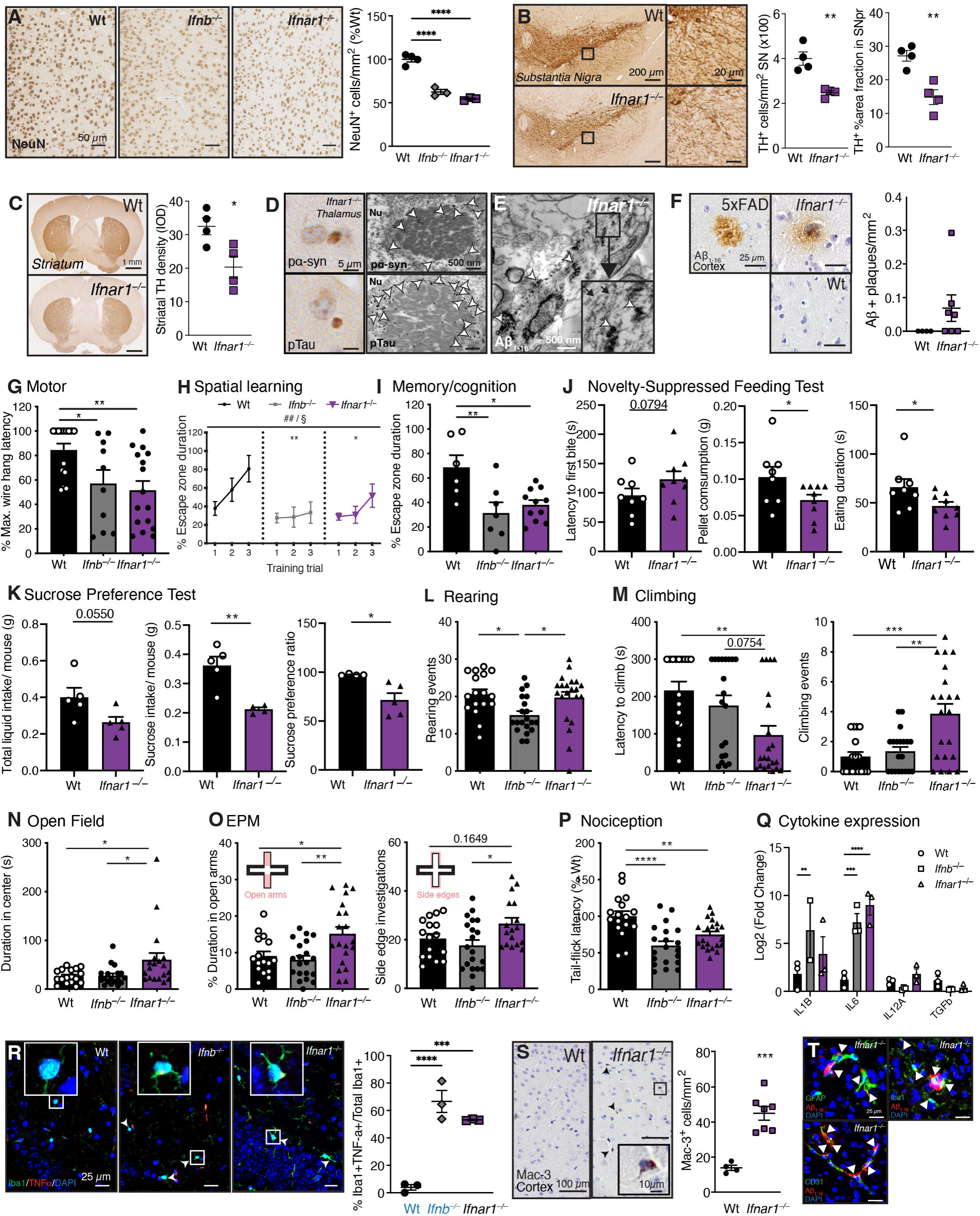
*Ifnar1^−/−^* leads to PDD-like neuropathology and behavior. (**A**) Immunohistochemical (IHC) images and collective quantification of NeuN^+^ cells per mm^2^ in 6-month-old Wt, *Ifnb^−/−^,* and *Ifnar1^−/−^* cortex, hippocampus, and olfactory bulb as %Wt (*n* = 3-4 per genotype). \*\*\*\**P* < 0.0001 by one-way ANOVA and Bonferroni’s post hoc correction test. Scale bars, 50 µm. (**B-C**) IHC images and quantifications of tyrosine hydroxylase (TH)^+^ DA neurons and neurites in 3-month-old Wt versus *Ifnar1^−/−^* (**B**) substantia nigra and (**C**) striatum. (*n* = 4 per genotype). \**P* < 0.05 and \*\**P* < 0.01 by unpaired *t*-test. Scale bars, 200 µm (substantia nigra), 20 µm (insert), and 1 mm (striatum). (**D)** IHC (left) and immunogold-labelled (white arrows) TEM images (right) for pα-syn or pTau of Lewy body (LB)-like inclusions in 12-month-old *Ifnar1^−/−^*thalamus. Nu = nucleus. Scale bars, 5 µm (IHC) and 500 nm (TEM). (**E**) TEM micrograph of immunogold-labelled Aβ_1-16_ (white arrows) in fibrillar structures (black arrows) in 12-mo *Ifnar1^−/−^* thalamus. (**F**) Representative images of 12-month-old 5xFAD, Wt, and *Ifnar1^−/−^*cortex stained for Aβ_1-16_, with extracellular Aβ^+^ plaques per mm^2^ cortex quantified in Wt versus *Ifnar1^−/−^*mice (*n* = 4-7 per genotype). Scale bars, 50 µm. **(G)** Wire hang performances of 3-month-old mice (*n* = 10-16 per genotype). \**P* < 0.05 and \*\**P* < 0.01 by one-way ANOVA and Tukey’s post hoc correction test. (**H-I**) Barnes maze results of 12-month-old mice (*n* = 6-10 per genotype). (**H**) Barnes maze spatial learning performances. ^##^*P* < 0.01 for training effect and ^§^P < 0.05 for genotype effect by two-way ANOVA. Genotype effect vs Wt \**P* < 0.05 and \*\**P* < 0.01 by Tukey’s post hoc test. (**I**) Memory probe on the Barnes maze as percent total test time spent in the escape zone. \**P* < 0.05 and \*\**P* < 0.01 by one-way ANOVA and Tukey’s post hoc correction test. (**J**) Novelty-suppressed feeding test results of 3-month-old mice (*n* = 8-9 per genotype), showing latency to first bite (seconds, s), amount of food consumed (grams, g), and eating duration (s). (**K**) Sucrose preference test results of 6-month-old Wt and *Ifnar1^−/−^* mice (*n* = 10-15 per genotype), comparing total liquid intake per mouse weight (g), sucrose intake per mouse weight (g), and sucrose preference ratio over water. \**P* < 0.05 and ** *P <* 0.01 by unpaired *t-*test. (**L**) Rearing activity of 3-month-old Wt, *Ifnb^−/−^,* and *Ifnar1^−/−^* mice (*n* = 17-20 per genotype). (**M**) Climbing activity of 3-month-old mice (*n* = 17-20 per genotype), quantified by latency to first climbing event (s) and number of climbing events. (**N**) Open field (OF) performance of 3-month-old mice as cumulative time (s) spent in maze center (*n* = 17-20 per genotype). (**O**) Elevated plus maze (EPM) performances of 3-month-old mice (*n* = 17-20 per genotype). Time spent in the maze open arms (top diagram, left graph) and number of side edge investigations (bottom diagram, right graph) were quantified. (**P**) Heat-induced tail flick latencies (nociception) of 3-month-old mice (*n* = 17-20 per genotype) as %Wt. (**Q**) Gene expression of cytokines IL-1b, IL-6, IL-12, and TGF-β by qPCR in 3-month-old mouse brain, shown as Log2FC of Wt (*n* = 3 per genotype). \*\**P* < 0.01, \*\*\**P* < 0.001, and \*\*\*\**P* < 0.0001 by two-way ANOVA and Tukey’s post hoc correction test. **(R**) IF images and quantification of TNF-ɑ^+^Iba1^+^ as %total Iba1^+^ microglia in the hippocampus of 9-month-old Wt, *Ifnb^−/−^,* and *Ifnar1^−/−^* mice (*n* = 3 per genotype). Previously published (12) Wt and *Ifnb^−/−^*data (blue text) were generated concurrently with the *Ifnar1^−/−^* data shown and included for comparison. \*\*\**P* < 0.001 and \*\*\*\**P* < 0.0001 by one-way ANOVA and Bonferroni’s post hoc correction test. Scale bars, 25 µm. (**S**) Representative IHC images and quantifications Mac-3^+^ microglia (black arrows) per mm^2^ in Wt vs *Ifnar1^−/−^* cortex (*n* = 4-7 per genotype). Scale bars, 100 µm; insert, 10 µm. \**P* < 0.05 by unpaired *t*-test. (**T**) IF images of GFAP^+^ astrocytes, Iba1^+^ microglia, or CD31^+^ endothelial cells (green) associated with Aβ^+^ (red) aggregates (white arrows) in 12-month-old *Ifnar1^−/−^* cortex. DAPI = blue. Scale bars, 25 µm. Data in all graphs are mean ±SEM.

To determine whether the identified neuropathologies resulted in discernible phenotypes, *Ifnar1^−/−^* mouse behaviors were assessed over time. 3-month-old *Ifnar1^−/−^* mice exhibited significantly reduced motor performances on the wire suspension test compared to Wt mice (Fig. 6G). Though 3-month-old *Ifnar1^−/−^* mice performed marginally better than *Ifnb^−/−^* mice on the Morris water maze (MWM) (Supplementary Fig. 5F-H), both 6-month-old *Ifnb^−/−^* mice and *Ifnar1^−/−^* mice demonstrated impaired spatial learning and memory on the Morris water maze (Supplementary Fig. 5I-K), which were sustained in 12-month-old *Ifnb^−/−^* and *Ifnar1^−/−^*mice on the Barnes maze (Fig. 6H, I).

Non-motor neuropsychiatric behaviors such as depression, anxiety, and pain, which can develop prior to PD diagnosis in humans (72, 73), were assessed in *Ifnar1^−/−^* mice. Depression-like behavior was observed on the novelty-suppressed feeding test, where 3-month-old *Ifnar1^−/−^* mice spent significantly less time eating and consumed less when exposed to a food pellet in a novel open arena after a 24 h fasting period (Fig. 6J). Similar behavior was sustained in 6-month-old *Ifnar1^−/−^*mice with the sucrose preference test, which manifested as significantly reduced sucrose preference ratio and reduced total sucrose intake compared to Wt mice (Fig. 6K). *Ifnb^−/−^* mice spontaneously reared less and for shorter durations than Wt and *Ifnar1^−/−^* mice (Fig. 6L; Supplementary Fig. 5L), corresponding with reduced motor capacity on the wire suspension test (Fig. 6G) and on the rotarod (12, 13); however, *Ifnar1^−/−^* mice initiated climbing behavior significantly faster and more frequently than Wt and *Ifnb^−/−^* mice (Fig. 6M). To determine whether *Ifnar1^−/−^* climbing behavior was due to increased anxiety, mouse behavior was assessed in open field (OF) and elevated plus maze (EPM). Genotypic differences were observed for distance travelled or velocity on either test (Supplementary Fig. 5M, N). In contrast with *Ifnb^−/−^* mice, which previously exhibited classic anxiety-like behaviors at a later age (Supplementary Fig. 5O), 3-month-old *Ifnar1^−/−^* mice spent more time in the OF center (Fig. 6N), on the EPM open arms, and had significantly more EPM side edge investigations than Wt and *Ifnb^−/−^*mice (Fig. 6O). Though conventionally interpreted as anxiolytic behavior, these data reflect similar escape-induced anxiety-like behavior on the EPM observed in the early stages of neurodegeneration in the 6-OHDA and A53T mouse models of PD (74, 75). Finally, both 3-month-old *Ifnb^−/−^* mice and *Ifnar1^−/−^* mice demonstrated faster heat-induced nociception, suggesting that IFNβ-IFNAR defects may be involved in early-onset pain sensitivity (Fig. 6P).

Lack of IFNβ-IFNAR signaling could impact expression of other brain cytokines like increased IL-1β, IL-6 and TNF-α (10, 12, 76, 77). To assess whether anxiety-related phenotypes could be associated with differential regulation of brain cytokines, a panel of pro- and anti-inflammatory cytokines were investigated by qPCR, among which IL-1β, IL-6, IL-12A and TGFβ were detectable (Fig. 6Q). Increased expression of proinflammatory IL-1β and IL-6 gene were found in cortex of mice lacking IFNβ-IFNAR signaling, however no significant differences were found between *Ifnb^−/−^* and *Ifnar1^−/−^*strains to correlate with differential anxiety-like behavior. Furthermore, as IFNβ loss was associated with elevated neuronal TNF-ɑ (12), *Ifnar1^−/−^* TNF-α^+^/Iba1^+^ counts were concurrently-quantified alongside Wt and *Ifnb^−/−^* counts (previously published (12), indicated in blue text). Like *Ifnb^−/−^* mice, higher numbers of TNF-α^+^ microglia in *Ifnar1^−/−^* mice were found compared to Wt controls (Fig. 6R), confirming neuroinflammation upon IFNβ-IFNAR loss. Significantly more phagocytic microglia, marked by lysosomal Mac-3 expression (78), were seen in *Ifnar1^−/−^*cortex compared to Wt (Fig. 6S). Interestingly, GFAP^+^ astrocytes and Iba1^+^ microglia were detected surrounding Aβ plaques (Fig. 6T, top panels), supporting astrocytic involvement in Aβ processing and microgliosis. Aβ was also found localized to CD31^+^ endothelial cells lining blood vessels in 12-month-old *Ifnar1^−/−^* cortex (Fig. 6T, bottom panel), mimicking what is observed in dementia (79, 80).

Collectively, these results show that molecular and functional differences observed due to IFNAR1 loss results in Parkinsonian dementia-like neuropathology, behavior deficits, and neuroinflammation.

### Neuronal and astrocytic IFNAR1 loss recapitulate distinct aspects of PDD-like neuropathology and behavior

As the transcriptomic and functional metabolic results suggested differential effects of IFNAR1 loss in neurons and astrocytes, PDD-like neuropathology and behavioral phenotypes were investigated in mice lacking either neuronal or astrocytic IFNAR1. Neuron-specific Cre recombinase (Cre)-floxed *Ifnar1* gene Syn1^Cre^;*Ifnar1^fl/fl^* mice and astrocyte-specific GFAP^Cre^;*Ifnar1^fl/fl^* mice were generated and validated for target-cell specific knock-down of *Ifnar1* using qPCR (Fig. 7A). Though the potential for GFAP^Cre^-based knockout to affect neuronal populations has been reported and discussed (81, 82), cell-specific deletion of *Ifnar1* in both astrocytes and neurons was confirmed, and no off-target reduction of *Ifnar1* were observed in either conditional mouse strains. Moreover, ISG15 downregulation was found in target cells but not in other brain cell types (Fig. 7B, C), supporting cell-specific loss of IFNAR signaling.

**Fig. 7.**
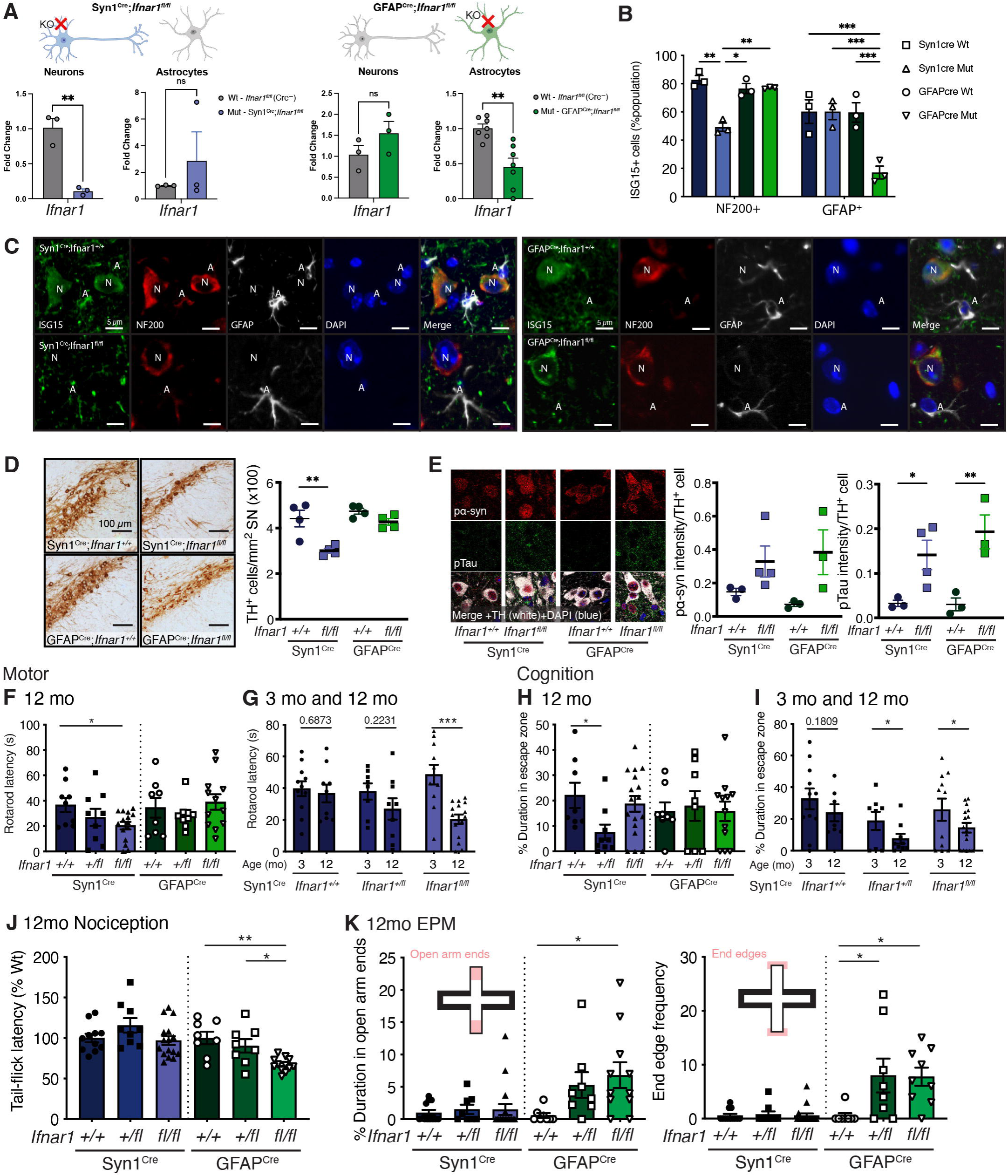
Neuron- or astrocyte-specific *Ifnar1^−/−^* recapitulate differential aspects of PDD-like phenotypes. (**A**) *Ifnar1* expression in primary cortical neurons vs enriched astrocyte cultures from Syn1^Cre^;*Ifnar1^fl/fl^* and GFAP^Cre^;*Ifnar1^fl/fl^*pups compared to Cre-negative *Ifnar1^fl/fl^* controls (*n* = 3-7 per genotype). \*\**P* < 0.01 by unpaired *t-*test. (**B-C**) Quantification (**B**) and Representative images (**C**) and of ISG15 (green) as %target population in 3-month-old Syn1Cre;*Ifnar1^+/+^*(Syn1cre Wt), Syn1Cre;*Ifnar1^fl/fl^* (Syn1cre Mut), GFAPCre;*Ifnar1^+/+^* (GFAPcre Wt), and GFAPCre;*Ifnar1^fl/fl^* (GFAPcre Mut) brains (n=3 per genotype). N = NF200^+^ neuron (red); A = GFAP^+^ astrocyte (white). DAPI = blue. Scale bars, 5 µm. *P < 0.05, **P < 0.01, and *** P < 0.001 by two-way ANOVA and Tukey’s post hoc test. (**D**) Representative images and quantifications of TH^+^ DA neurons and neurites in substantia nigra of 3-month-old Syn1^Cre^;*Ifnar1*^fl/fl^ mice and astrocytic GFAP^Cre^;*Ifnar1*^fl/fl^ mice compared to littermate controls (*n* = 3-4 per genotype). \*\**P* < 0.01 by one-way ANOVA and Bonferroni’s post hoc correction test. Scale bars, 100 µm. (**E**) Representative images and quantifications of pα-syn (red) and pTau (green) fluorescence intensity per TH^+^ (white) neurons of 3-month-old Syn1^Cre^;*Ifnar1*^fl/fl^ mice and astrocytic GFAP^Cre^;*Ifnar1*^fl/fl^ mice compared to littermate controls (*n* = 3-4 per genotype). Blue = DAPI. \**P* < 0.05 and \*\**P* < 0.01 by one-way ANOVA and Bonferroni’s post hoc correction test. Scale bars, 100 µm. (**F**) Rotarod performance of 12-month-old neuronal Syn1^Cre^;*Ifnar1*^fl/fl^ mice and astrocytic GFAP^Cre^;*Ifnar1*^fl/fl^ mice (*n* = 7-16 per genotype). \**P* < 0.05 by one-way ANOVA and Tukey’s post hoc correction test. (**G**) Rotarod comparison of 3-month-old (*n* = 8-11 per genotype) and 12-month-old (*n* = 7-16 per genotype) neuronal Syn1^Cre^;*Ifnar1*^fl/fl^ mice. \*\*\**P* < 0.001 by unpaired *t-*test. (**H**) Barnes maze probe performance reflecting cognitive capacity of 12-month-old neuronal Syn1^Cre^;*Ifnar1*^fl/fl^ mice and astrocytic GFAP^Cre^;*Ifnar1*^fl/fl^ mice (*n* = 7-16 per genotype). \**P* < 0.05 by one-way ANOVA and Tukey’s post hoc correction test. **(I)** Barnes maze probe performance comparison between 3-month-old (*n* = 8-11 per genotype) and 12-month-old (*n* = 7-16 per genotype) neuronal Syn1^Cre^;*Ifnar1*^fl/fl^ mice. \**P* < 0.05 by unpaired *t-*test. Data in all graphs are mean ±SEM. (**J**) Tail flick latencies representing heat-induced nociception as %Wt (either Syn1^Cre^;*Ifnar1*^+/+^ or GFAP^Cre^;*Ifnar1^+/+^*, respectively) of 3-month-old (*n* = 8-11 per genotype) and 12-month-old mice (*n* = 7-16 per genotype). (**K**) EPM end-edge investigations of 12-month-old Syn1^Cre^;*Ifnar1^fl/fl^* and GFAP^Cre^;*Ifnar1^fl/fl^* mice (*n* = 7-16 per genotype). \**P* < 0.05 by one-way ANOVA and Tukey’s post hoc correction test. Data in all graphs are mean±SEM.

As Cre expression can in itself alter anxiety-related behaviors (83), Syn1^Cre^;*Ifnar1^+/+^*and GFAP^Cre^;*Ifnar1^+/+^* littermates were used as controls for both molecular and behavioral comparisons. Significant DA neuronal loss in the substantia nigra was observed specifically in 3-month-old neuronal Syn1^Cre^;*Ifnar1^fl/fl^*mice but not in astrocytic GFAP^Cre^;*Ifnar1^fl/fl^* mice or Syn1^Cre^;*Ifnar1^+/+^* littermate controls (Fig. 7D). Accumulation of pα-syn and pTau was found in TH^+^ neurons of both Syn1^Cre^;*Ifnar1^fl/fl^* and GFAP^Cre^;*Ifnar1^fl/fl^*mice (Fig. 7E); however, only neuronal Syn1^Cre^;*Ifnar1^fl/fl^*mice had indications of motor and cognitive deficits at 3 months of age (Supplementary Fig. 6A, B), which became fully penetrant at 12 months of age (Fig. 7F-I) as shown by significant motor impairment on the rotarod (Fig. 7F, G) and reduced cognitive performance on the Barnes maze (Fig. 7H, I). Though cognitive deficits were only significant in 12-month-old neuronal heterozygous Syn1^Cre^;*Ifnar1^+/fl^* mice (Fig. 7H), both heterozygous and homozygous groups showed significantly inhibited Barnes maze performances compared to their respective 3-month-old groups (Fig. 7I). Aging differences were not significant between 3- and 12-month-old control Syn1^Cre^;*Ifnar1^+/+^*mice (Fig. 7I). Of note, neither astrocytic GFAP^Cre^;*Ifnar1^+/fl^* nor GFAP^Cre^;*Ifnar1^fl/fl^* demonstrated motor deficits or cognitive decline at either time point (Fig. 7F, H; Supplementary Fig 6A, B).

Behaviors resembling neuropsychiatric abnormalities, which manifested prior to cognitive decline in *Ifnar1^−/−^*mice, were assessed in Syn1^Cre^;*Ifnar1^fl/fl^* mice and GFAP^Cre^;*Ifnar1^fl/fl^* mice. Though pain sensitivity was not detected in 3-month-old conditional knockout animals (Supplementary Fig. 6C), significantly increased heat-induced nociception manifested only in 12-month-old GFAP^Cre^;*Ifnar1^fl/fl^* mice (Fig. 7J). Like *Ifnar1^−/−^* mice, hyper-anxious behavior was significant in 3-month-old GFAP^Cre^;*Ifnar1^fl/fl^* mice (Supplementary Fig. 6D) and more pronounced in 12-month-old GFAP^Cre^;*Ifnar1^+/fl^*mice and GFAP^Cre^;*Ifnar1^fl/fl^* compared to GFAP^Cre^;*Ifnar1^+/+^*controls or Syn1^Cre^;*Ifnar1^fl/fl^* mice (Fig. 7K; Supplementary Fig. 6E). The absence of neuropsychiatric phenotypes in neuronal Syn1^Cre^;*Ifnar1^fl/fl^* mice, while being exhibited by GFAP^Cre^;*Ifnar1^fl/fl^* mice, suggests a unique role for astrocytic IFNAR1 in preventing the development of neuropsychiatric symptoms associated with PDD.

In summary, neuronal IFNAR1 loss appeared sufficient to recapitulate substantia nigral neuropathology alongside motor and cognitive behavior deficits, whereas astrocytic IFNAR1 loss recapitulated behavior resembling neuropsychiatric abnormalities. The later-onset of behavior deficits in both cell-specific IFNAR1 knock-out strains relative to genomic *Ifnar1^−/−^*mice suggest that synergistic dysfunction occurs upon neuronal and astrocytic IFNAR1 loss, contributing to a progressive PDD-like phenotype in *Ifnar1^−/−^*mice.

This study shows that cell-specific transcriptional alterations in mitochondrial and energy metabolism genes in *Ifnar1^−/−^* brain are reflected in regional proteome changes and altered glucose metabolism, supporting a regulatory role of IFNAR1 in excitatory neurotransmission. These molecular changes upon IFNAR1 loss results in broad neurodegenerative processes as well as neuropathology resembling PDD in aged mice, including LB-like inclusions, pTau accumulation, Aβ pathology, gliosis with neuroinflammatory profile associated with mitochondrial dysfunctions. Importantly, neuronal IFNAR1 loss was sufficient to induce PD-like neurodegeneration and motor and cognitive behavior manifestations, whereas astrocytic IFNAR1 loss manifested as behaviors resembling pain sensitivity and hyper-anxiety, together revealing distinct brain cell-specific requirements of IFNAR1 signaling which in concert contribute to maintaining brain homeostasis (Fig. 8).

**Fig. 8.**
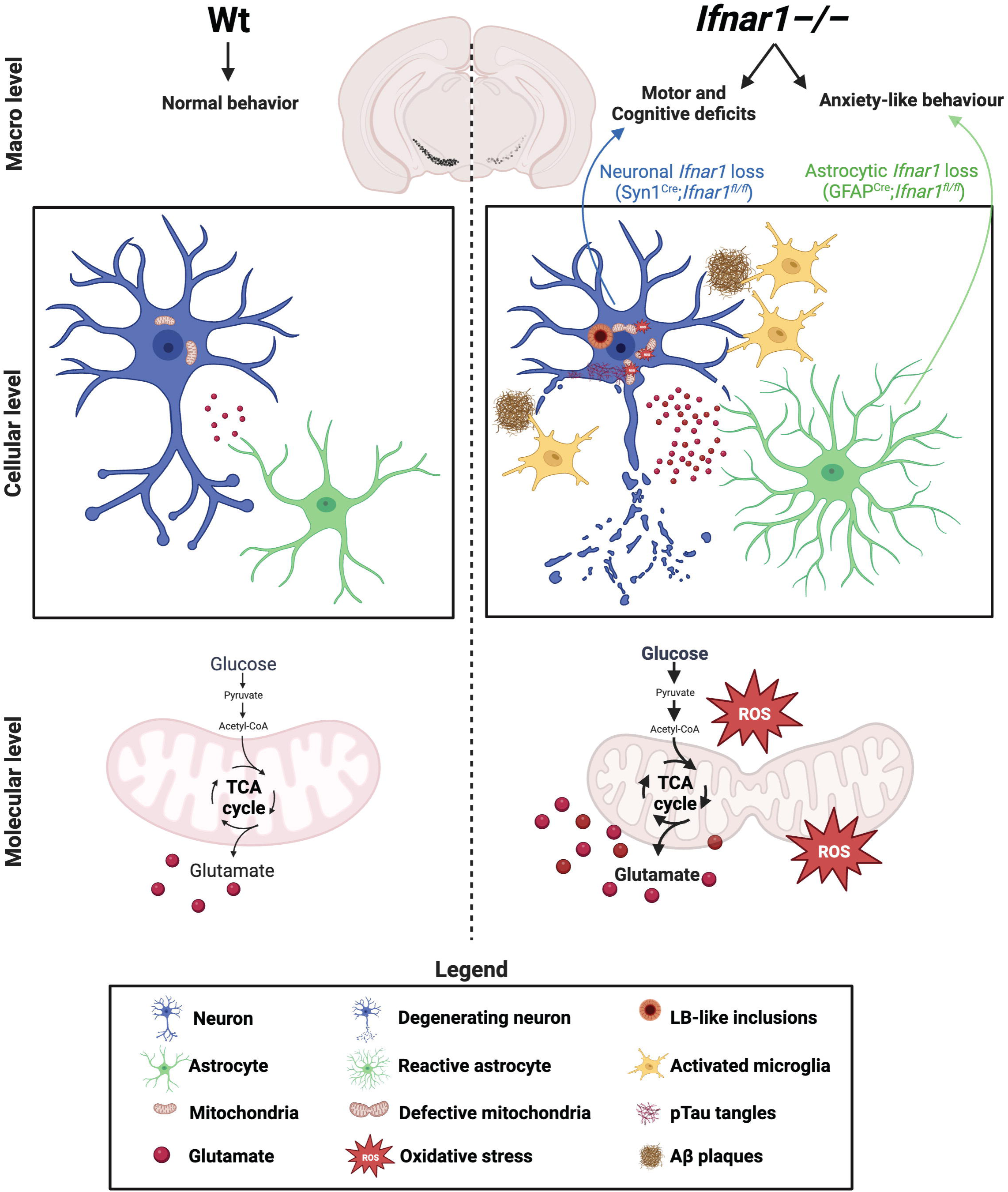
Graphical summary of the study. Lack of immune modulation due to IFNAR1 loss results in mitochondrial dysfunction shown as mitophagy deficits, oxidative stress, and glucose hypermetabolism (molecular level), leading to development of neuroinflammation, neurodegeneration, LB-like inclusions, pTau accumulation, Aβ plaques, and disturbed brain energy metabolism (cellular level). These PDD-like pathologies manifest as motor, cognitive, and neuropsychiatric behavior deficits upon aging (macro level). Cardinal motor and cognitive manifestations were recapitulated by mice lacking neuronal IFNAR1, and neuropsychiatric disturbance-like behaviors were driven by astrocytic IFNAR1 loss. Therapeutics targeting the synergistic dysfunction that results from neuronal and astrocytic IFNAR1 loss may therefore prevent PDD progression and manifestation.

## Discussion

IFNβ is a potent anti-inflammatory and antiviral cytokine that signals through IFNAR1 and IFNAR2 receptors, and defective signaling of IFNβ through IFNAR is associated with patients suffering from PD and PDD (10) and mimics PDD-like pathology in experimental models (11–13). However, the neuropathological and neuropsychiatric outcomes resulting from *Ifnb* loss may differ in some respects from those caused by the absence of the *Ifnar1* gene, which could be due to potential compensatory mechanisms involving other type I interferon cytokines or variations related to the cellular origins of the defective genes. This report explores the role of brain cell-specific IFNAR1 and provides evidence of the importance of IFNAR1, conventionally investigated in immune cells (84) or upon viral infection (85–87), in neuronal homeostasis, including how its dysfunction contributes to neuropathological and neuropsychiatric conditions that resemble PDD. Baseline IFNAR1 expression differs in neurons and astrocytes and is distinctly impacted in LBD and PD along with several related Type I-IFN genes, implicating dysregulated IFNAR1 signaling in dementia development as previously observed in sporadic PDD (10). Unbiased transcriptomic and proteomic analysis revealed molecular differences affecting pathways related to mitochondrial function and PD, manifesting as brain metabolic dysregulation and clinical and pathological phenotypes resembling PDD in *Ifnar1^−/−^* mice. These findings support that lack of IFNβ-IFNAR signaling in *Ifnar1^−/−^* mice, like in *Ifnb^−/−^* mice, leads to dysregulated neuronal autophagy and mitophagy (11–13, 88) and immunomodulation (12, 17, 50), resulting in neuroinflammation alongside mitochondrial and metabolic malfunctions (10, 52). Importantly, it is shown that lack of neuronal IFNAR1 (Syn1^Cre^;*Ifnar1^fl/fl^*) alone mimics all *Ifnar1^−/−^* behavioral manifestations except for behavior resembling neuropsychiatric disturbances and pain sensitivity, which manifested instead in mice lacking astrocytic IFNAR1 (GFAP^Cre^;*Ifnar1^fl/fl^*), revealing distinct cell-specific requirements of IFNAR1 in brain homeostasis.

Collectively, the transcriptomic, proteomic, and functional investigations in this report suggest that excitatory neurotransmission is compromised in *Ifnar1^−/−^* brains, and is supported by another study implicating IFNAR1 loss in the dysregulation of the glutamate-aspartate transporter GLAST and subsequent detrimental effects on synaptic strength (14). Consistent with previous work showing that IFNβ signaling is essential for mitochondrial homeostasis (52), the transcriptomic and proteomic results here suggested dysregulated processes essential for neurotransmission upon IFNAR1 loss, including mitochondrial function (63, 64). Additionally, we show here that lack of *Ifnar1*, like in *Ifnb^−/−^* mice, causes accumulation of defective/oxidatively damaged mitochondria. Together, these findings support the requirement of IFNβ-IFNAR signaling in mitochondrial homeostasis (10, 11, 52) and shed further light on related reports in PDD patients implicating mitochondrial dysfunction in dementia development (10, 89, 90).

Type-I IFNs have been shown to increase fatty acid oxidation and oxidative phosphorylation in peripheral immune cells (91) and it was found that IFNβ regulates cellular glucose metabolism upon viral infection (92); however, the investigation of IFNAR1 involvement in brain energy metabolism is unprecedented. Brain glucose metabolism is closely coupled to glutamatergic neurotransmission (69, 93), which was found here to be hypermetabolic in *Ifnar1^−/−^* cortex and hippocampus. Increased glucose metabolism has been reported prior to proteinopathy in AD mouse models (41, 94) and correlated to cognitive dysfunction severity in PD patients (7). Brain glucose hypermetabolism upon IFNAR1 loss may therefore reflect compensatory metabolic excitatory mechanisms for declining neurotransmission that manifests upon aging as PDD-like behaviors in *Ifnar1^−/−^* mice.

The neurodegeneration observed in *Ifnar1^−/−^* mice, as with *Ifnb^−/−^* mice (12, 13), is comprised of significant DA neuron loss, the hallmark of PD pathology, alongside cortical and hippocampal neurodegeneration mimicking cortical thinning and hippocampal atrophy among PDD patients (24, 95). Though DA neurodegeneration in the substantia nigra is significant in PD patients and considered a defining neuropathology of parkinsonism (96), PD neuropathology extends throughout the brain beyond the basal ganglia (24, 95), and the impact of L-dopa treatment, which targets DA neuronal dysfunction, on prevention of non-motor symptoms is poor (97–99). Thus, the broad behavioral phenotypes exhibited by mice lacking IFNAR1 signaling, which appear to phenocopy patients with PD/PDD, likely reflect a general failure in neuronal health and signaling abilities and highlight a need for investigating neurodegenerative mechanisms beyond DA neuropathology in PDD. Though the mechanistic observations in this report were made in mice, patient-based studies showing a reduced risk of PD development in hepatitis C patients receiving IFN-I treatment (100) as well as significant reduction of IFNAR1 in PDD compared to PD patients without dementia (10) support the importance of the homeostatic functions of IFNAR1 in maintenance of human brain health.

Lack of astrocytic IFNAR1 signaling may contribute to neuropsychiatric abnormalities in PDD, including depression, early-onset anxiety and pain sensitivity, due to its impact on hippocampal metabolic regulation reported here to be disturbed in *Ifnar1^−/−^* mice. Moreover, the earlier and more pronounced manifestation of anxiety-like behavior of *Ifnar1^−/−^*compared to *Ifnb^−/−^* mice indicate that although IFNβ and IFNAR1 work together to conduct cellular signaling, their function or lack of it do not necessarily exert the exact same effect. Depending on the cellular source of IFNβ production, versus cells actively expressing IFNAR1, there might be differences in outcome upon autocrine and/or paracrine functions. In support, neuron-derived IFNβ, but not astrocytic or microglial-derived IFNβ, can exclusively convert encephalitogenic T cells into regulatory T cells in a neuroinflammatory context by autocrine regulation of neuronal PD-L1 expression and negative PD-1 dependent T cell suppression (50) and glioblastoma control (101). Additionally, residual IFNɑ-IFNAR signaling in *Ifnb^−/−^*mice could potentially exert functions distinct from IFNβ-IFNAR (102), particularly in astrocytes versus neurons compared to complete lack of signaling or differential signaling in *Ifnar1^−/−^* brain, thus leading to different outcome.

Though cognitive deficiencies were previously reported in young mice lacking astrocytic IFNAR1 (1-1.5-month-old) GFAP^Cre^;*Ifnar^+/fl^*mice compared to Cre-negative (*Ifnar1^fl/fl^*) mice on the Morris Water Maze (14), here we did not observe cognitive deficits at any age between 3-12-month-old GFAP^Cre^;*Ifnar^+/fl^* and GFAP^Cre^ controls on the Barnes maze here, which could be due to differences in age, test conditions, and the use of Cre+ in our study versus Cre-mice as controls in the previous report (14). Furthermore, though cell-specific IFNAR1 loss was confirmed in the conditional strains both here and previously (85), more recent literature have reported some limitations in usage of the GFAP^Cre^ lines which are associated either with increased toxicity of some Cre lines or lack of efficient gene editing because of lower expression of GFAP in a subtype of astrocytes during adulthood in striatum (103, 104), as well as their potential to affect distinct neuronal populations (81, 82). A less pronounced cognitive deficit observed in homozygous versus heterozygous Syn1^Cre^ animals may indicate activation of compensatory signaling in astrocytes or other supportive brain cell types. Nevertheless, the delayed onset of behavior deficits in either cell-specific knockout strain indicates a synergistically detrimental effect of IFNAR1 loss in both neurons and astrocytes that results in PDD-like neuropathology and behavioral phenotypes. Further study of brain cell-specific roles of IFNβ-IFNAR signaling will prove essential in understanding neurodegenerative pathophysiology and identifying more precise therapeutic targets.

Together, these findings provide molecular insight into cell-specific roles of IFNAR1 in brain homeostasis, loss of which results in alterations of genes and proteins indicative of lack of neuronal immunomodulation, neuroinflammation, mitochondrial defects, and dysregulated brain energy metabolism prior to the development of PDD-like neurodegeneration, proteinopathy, and behavior abnormalities. Importantly, neuron-specific IFNAR1 loss alone induced the development of PDD-like neurodegeneration and behavioral abnormalities, highlighting the importance of functional neuronal IFNAR1 in brain homeostasis. These results also suggest a significant role for astrocytic IFNAR1 in neuronal support and modulation of neuropsychiatric-like outcomes. As dysfunctional IFNβ-IFNAR signaling has been reported to be associated with human sporadic PD and its progression to PDD (10), results from this study suggest that concerted and targeted repair of dysfunctional neuronal and astrocytic IFNAR1 may be a promising therapeutic strategy in mitigating different aspects of disease associated with dementia development, particularly in PDD.

## Acknowledgements

This work was supported by grants from the Lundbeck Foundation (E.B.V. PhD fellowship R191-2015-1605 and project support to S.I.-N. R223-2016-849 and LF Professorship grant R359-2020-2658), the BRIDGE Translational Excellence Programme (bridge.ku.dk) at the Faculty of Health and Medical Sciences, University of Copenhagen, funded by the Novo Nordisk Foundation (E.B.V. grant agreement NNF20SA0064340), the EU Horizon 2020 Research and Innovation Programme (Z.Z. Marie Sklodowska-Curie DISCOVER grant agreement 10103491), and the Danish Council for Independent Research-Medicine (S.I.-N. grants DFF-6110-00658 and DFF-2034-00015B). Mass spectrometry experiments were conducted at the Novo Nordisk Foundation Centre for Protein Research (CPR, NNF14CC0001). The authors declare no competing financial interests.

The authors thank Prof. Konstantin Khodosevich and Dr. Mykhailo Batiuk for guidance with snRNA-seq analysis, Natasha Fauerby for technical support, Joana Marturià-Navarro for help with primary neuronal cultures, Dr. Yawei Liu for FACS and qPCR expertise, Dr. Ana Rita Freitas Colaço for proteomics analysis coding support, Dr. Leonora Rib for helping combine the transcriptomic and proteomic datasets, and Dr. Neža Cankar for proofreading the manuscript. Schematics were generated using Biorender.com.

## Author contributions

E.B.V., Z.Z., J.L., J.V.A., F.L.Q., E.W.W., A.M., G.J.-D., L.R.-P., E.T., O.K., D.L., T.G., T.B., M.P., B.I.A., M.M., and N.H.S. generated data for the manuscript. E.B.V collected brain tissues for immunoblots, snRNA-seq, and LC-MS/MS. Z.Z. performed neuronal and astrocytic primary cultures, qPCR, FC, GSEA of transcriptomic data, and combined visualization of transcriptomic and proteomic datasets. J.L. conducted snRNA-seq experiments, transcriptomic data processing, analysis, and visualization. F.L.Q., M.M., and N.H.S. conducted LC-MS/MS experiments, proteomic data processing, analysis, and visualization. J.V.A., E.W.W., and B.I.A. performed acute brain slice incubations, GC-MS, HPLC, and analysis. E.B.V. and A.M. conducted behavior experiments and analysis. L.R.-P. conducted immunoblots and analysis. E.T., O.K., and D.L. conducted electron microscopy experiments. E.B.V, A.M., G.J.-D., T.G., T.B., and M.P. conducted immunohistochemical experiments and analysis. E.B.V. and S.I.-N. conceived the project and wrote the manuscript. All authors reviewed and edited the manuscript.

## Ethics

All animal experiments were performed in accordance with Danish ethical standards in compliance with Directive 2010/63/EU, under ethical permission licenses 2013-15-2934-00807 and 2018-15-0201-01572, and are in adherence with ARRIVE guidelines (v2.0).

## Data Availability

The snRNA-seq data has been deposited in the Gene Expression Omnibus (GEO) with the identifier GSE213671 (https://www.ncbi.nlm.nih.gov/geo/query/acc.cgi?acc=GSE213671).

MS-based proteomics data is deposited to the ProteomeXchange Consortium (http://proteomecentral.proteomexchange.org) via the PRIDE partner repository(105) with the identifier PXD037113 (http://www.ebi.ac.uk/pride/archive/projects/PXD037113).

## Consent to Participate and Consent to Publish declarations

Not applicable.

## Supplementary Figure Legends

**Supplementary Fig. 1. Additional snRNA-seq analysis supplementing Fig. 2.** (**A**) Uniform manifold projections (UMAPs) showing complete annotations of all cell types and sub-types defining the broad cell classes used for DE analysis. (**B**) Unbiased hierarchical clustering of differentially expressed genes among all 5 cortical cell classes isolated from 1.5-month-old Wt and *Ifnar1^−/−^* mice showing similarities between samples. (**C**) GSEA pathway enrichment scores for pathways relating to interferon signaling, inflammation, and cytokine signaling in Glutamatergic and GABAergic cell classes. NES = Normalized Enrichment Score. (**D**) Relative mRNA expression of IFNAR1 and ISG15 expression in the cortex of 3-month-old Wt, *Ifnb^−/−^*, and *Ifnar1^−/−^* mice (*n* = 3 per genotype) as log2FC expression standardized to two housekeeping genes (*Beta-Actin* and *Rpl13a*) and normalized to Wt. \**P* < 0.05 by two-way ANOVA and Tukey’s post hoc test. (**E**) Quantification of parent (%) frequency of ISG15-PE single cells in NF200-FITC neuronal, GLAST-APC astrocyte, and ‘All’ single cell populations analyzed by flow cytometry in 3-month-old Wt and *Ifnar1^−/−^* cortex, normalized to Wt (*n* = 4 per genotype). \**P* < 0.05 by two-way ANOVA and Bonferroni’s post hoc test. (**F-G**) GSEA pathway tables showing top 15 positively and negatively enriched terms for (**F**) microglia (Reactome and KEGG) and (**G**) oligodendrocytes (Reactome). Significant neurodegenerative disease pathways are highlighted in bold text.

**Supplementary Fig. 2. Dysregulated GSEA pathways highlighting Parkinson’s Disease and mitochondrial metabolism pathways in *Ifnar1^−/−^* neuronal sub-types and astrocytes.** (**A-B**) Top 20 upregulated KEGG pathways cortical (**A**) glutamatergic and (**B**) GABAergic neuronal subtypes annotated in the snRNA-seq dataset shown in Supplementary Fig. 1B, highlighting ‘Parkinsons disease’ (green boxes), ‘oxidative phosphorylation’ (orange boxes), and other neurodegenerative disease pathways such as ‘Alzheimers disease’ and ‘Huntingtons disease’ (purple boxes). (**C**) GSEA pathway enrichment scores for pathways relating to energy metabolism and mitochondrial function in cortical *Ifnar1^−/−^* vs Wt glutamatergic and GABAergic neurons. NES = normalized enrichment score. (**D-E**) Heatmaps showing differential gene expression within the major cortical cell classes of genes within the Reactome pathways (**E**) ‘Trafficking of GluR2 containing AMPA receptors’ and (**E**) **‘**Highly calcium permeable nicotinic acetylcholine receptors’ identified as commonly affected in *Ifnar1^−/−^* glutamatergic and GABAergic neuronal classes shown in Fig. 4L and M, respectively. (**F**) GSEA pathway enrichment scores for pathways relating to neurotransmission and inflammation in cortical *Ifnar1^−/−^* vs Wt astrocytes. NES = normalized enrichment score. (**G**) showing differential gene expression among the major cortical cell classes of genes within the Reactome pathway ‘Pyruvate metabolism’ identified in *Ifnar1^−/−^* astrocytes shown in Fig. 2N.

**Supplementary Fig. 3. Additional LC-MS/MS proteomics analysis supplementing Fig. 3.** (**A-C**) Quality control for proteomics analysis. (**A**) Total number of proteins per sample. (**B**) Coefficients of variation (CVs) between samples. (**C**) Protein intensities, showing similarity between samples.

**Supplementary Fig. 4. Additional metabolic isotope labelling data supplementing Fig. 5** (**A-B**) Cortex and hippocampal slices from 3-month-old Wt and *Ifnar1^−/−^* mice (*n* = 6-7 per genotype) were incubated with (**A**) [U-^13^C]glutamine or (**B**) [U-^13^C]glutamate, which primarily reflect neuronal and astrocytic metabolism, respectively. \**P* < 0.05 by *t-*test.

**Supplementary Fig. 5. Additional neuropathological and behavior data in *Ifnar1^−/−^*supplementing Fig. 6.** (**A-C**) Representative images and quantification of NeuN+ cells in (**A**) cortex, (**B**) hippocampus, and (**C**) olfactory bulb of 6-month-old Wt, *Ifnb^−/−^*, and *Ifnar1^−/−^* mice as %Wt (*n* = 3-4 per genotype). \**P* < 0.05 by one-way ANOVA and Dunnett’s post hoc correction test. (**D**) Representative images and quantification of %loss of TH^+^ cells in the substantia nigra pars compacta (SNc) versus ventral tegmental area (VTA) of 3-month-old *Ifnar1^−/−^* mice compared to Wt mice (*n* = 4 per genotype). \*\**P* < 0.01 by *t*-test. (**E**) Representative immunoblots and quantifications of relative protein levels of total tau (Tau pan), phosphorylated tau at threonine 205 (pTau), alpha-synuclein (α-syn), and phosphorylated α-syn at serine 129 (pα-syn) in brain lysates from 1.5- and 12-month-old Wt vs *Ifnar1^−/−^* mice (*n* = 4 per genotype), normalized to vinculin. \**P* < 0.05 by *t*-test. (**F-K**) MWM results for 3-month-old **(F-H)** and 6-month-old **(I-K)** Wt, *Ifnb^−/−^*, and *Ifnar1^−/−^* mice. Data are mean ±SEM, *n* = 13-20 per genotype. (**F, I**) MWM training performances. ^§§^*P* < 0.01 for genotype effect by two-way ANOVA. Genotype effect vs Wt \**P* < 0.05 and \*\**P* < 0.01 by Tukey’s post hoc test. (**G, J**) Time spent in platform zone as % duration of test session. \**P* < 0.05 and \*\**P* < 0.01 by one-way ANOVA and Tukey’s post hoc correction test. (**H, K**) Number of platform zone crossings. \**P* < 0.05 by one-way ANOVA and Tukey’s post hoc correction test. (**L**) Additional rearing activity measurements of 3-month-old Wt and *Ifnar1^−/−^* mice (*n* = 10-15 per genotype), including latency to begin rearing (s) and cumulative rearing duration (s). (**M**) Distance (cm) and velocity (cm/s) of 3-month-old Wt, *Ifnb^−/−^*, and *Ifnar1^−/−^* mice (*n* = 17-20 per genotype) on the OF, corresponding to Fig. 6Q. (**N**) Distance and velocity of 3-month-old mice on the EPM, corresponding to Fig. 6R. (**O**) EPM Open arm duration and side edge investigations of 6-month-old mice (*n* = 10-17 per genotype).

**Supplementary Fig. 6. Additional data on conditional *Ifnar1^−/−^* strains supplementing Fig. 7.** (**A-B**) Rotarod (**A**) and Barnes Maze (**B**) performances of 3-month-old neuronal Syn1^Cre^;*Ifnar1^fl/fl^* mice and astrocytic GFAP^Cre^;*Ifnar1^fl/fl^* mice (*n* = 8-11 per genotype). (**C**) Nociception to heat-induced pain as %Wt performance (either Syn1^Cre^;*Ifnar1*^+/+^ or GFAP^Cre^;*Ifnar1^+/+^*, respectively) of 3-month-old Syn1^Cre^;*Ifnar1^fl/fl^* and GFAP^Cre^;*Ifnar1^fl/fl^* mice (*n* = 8-11 per genotype). (**D-E**) EPM performances of 3-month-old (**D**) Syn1^Cre^;*Ifnar1^fl/fl^* and GFAP^Cre^;*Ifnar1^fl/fl^*mice (*n* = 8-11 per genotype) and 12-month-old (**E**) Syn1^Cre^;*Ifnar1^fl/fl^*and GFAP^Cre^;*Ifnar1^fl/fl^* mice (*n* = 7-16 per genotype). \**P* < 0.05 by one-way ANOVA and Tukey’s post hoc correction test. Data in all graphs are mean ±SEM.

## References

1. Hely MA, Reid WG, Adena MA, Halliday GM, Morris JG. The Sydney multicenter study of Parkinson’s disease: the inevitability of dementia at 20 years. Mov Disord. 2008;23(6):837–44.

2. Petrou M, Bohnen NI, Müller ML, Koeppe RA, Albin RL, Frey KA. Aβ-amyloid deposition in patients with Parkinson disease at risk for development of dementia. Neurology. 2012;79(11):1161–7.

3. Shah N, Frey KA, Müller ML, Petrou M, Kotagal V, Koeppe RA, et al. Striatal and Cortical β-Amyloidopathy and Cognition in Parkinson’s Disease. Mov Disord. 2016;31(1):111–7.

4. Ferreira D, Przybelski SA, Lesnick TG, Lemstra AW, Londos E, Blanc F, et al. β-Amyloid and tau biomarkers and clinical phenotype in dementia with Lewy bodies. Neurology. 2020;95(24):e3257–e68.

5. Peppard RF, Martin WRW, Carr GD, Grochowski E, Schulzer M, Guttman M, et al. Cerebral Glucose Metabolism in Parkinson’s Disease With and Without Dementia. Archives of Neurology. 1992;49(12):1262–8.

6. Dunn L, Allen GF, Mamais A, Ling H, Li A, Duberley KE, et al. Dysregulation of glucose metabolism is an early event in sporadic Parkinson’s disease. Neurobiology of aging. 2014;35(5):1111–5.

7. Meles SK, Renken RJ, Pagani M, Teune LK, Arnaldi D, Morbelli S, et al. Abnormal pattern of brain glucose metabolism in Parkinson’s disease: replication in three European cohorts. European Journal of Nuclear Medicine and Molecular Imaging. 2020;47(2):437–50.

8. Blum D, la Fougère C, Pilotto A, Maetzler W, Berg D, Reimold M, et al. Hypermetabolism in the cerebellum and brainstem and cortical hypometabolism are independently associated with cognitive impairment in Parkinson’s disease. Eur J Nucl Med Mol Imaging. 2018;45(13):2387–95.

9. Gordon BA, Blazey TM, Su Y, Hari-Raj A, Dincer A, Flores S, et al. Spatial patterns of neuroimaging biomarker change in individuals from families with autosomal dominant Alzheimer’s disease: a longitudinal study. The Lancet Neurology. 2018;17(3):241–50.

10. Magalhaes J, Tresse E, Ejlerskov P, Hu E, Liu Y, Marin A, et al. PIAS2-mediated blockade of IFN-β signaling: a basis for sporadic Parkinson disease dementia. Molecular Psychiatry. 2021.

11. Tresse E, Marturia-Navarro J, Sew WQG, Cisquella-Serra M, Jaberi E, Riera-Ponsati L, et al. Mitochondrial DNA damage triggers spread of Parkinson’s disease-like pathology. Molecular Psychiatry. 2023.

12. Villanueva EB, Tresse E, Liu Y, Duarte JN, Jimenez-Duran G, Ejlerskov P, et al. Neuronal TNFα, Not α-Syn, Underlies PDD-Like Disease Progression in IFNβ-KO Mice. Ann Neurol. 2021;90(5):789–807.

13. Ejlerskov P, Hultberg JG, Wang J, Carlsson R, Ambjorn M, Kuss M, et al. Lack of Neuronal IFN-beta-IFNAR Causes Lewy Body- and Parkinson’s Disease-like Dementia. Cell. 2015;163(2):324–39.

14. Hosseini S, Michaelsen-Preusse K, Grigoryan G, Chhatbar C, Kalinke U, Korte M. Type I Interferon Receptor Signaling in Astrocytes Regulates Hippocampal Synaptic Plasticity and Cognitive Function of the Healthy CNS. Cell reports. 2020;31(7):107666.

15. Bakken TE, Jorstad NL, Hu Q, Lake BB, Tian W, Kalmbach BE, et al. Comparative cellular analysis of motor cortex in human, marmoset and mouse. Nature. 2021;598(7879):111-9.

16. Kamath T, Abdulraouf A, Burris SJ, Langlieb J, Gazestani V, Nadaf NM, et al. Single-cell genomic profiling of human dopamine neurons identifies a population that selectively degenerates in Parkinson’s disease. Nature Neuroscience. 2022;25(5):588–95.

17. Prinz M, Schmidt H, Mildner A, Knobeloch K-P, Hanisch U-K, Raasch J, et al. Distinct and Nonredundant In Vivo Functions of IFNAR on Myeloid Cells Limit Autoimmunity in the Central Nervous System. Immunity. 2008;28(5):675–86.

18. Kamphuis E, Junt T, Waibler Z, Forster R, Kalinke U. Type I interferons directly regulate lymphocyte recirculation and cause transient blood lymphopenia. Blood. 2006;108(10):3253–61.

19. Zhu Y, Romero MI, Ghosh P, Ye Z, Charnay P, Rushing EJ, et al. Ablation of NF1 function in neurons induces abnormal development of cerebral cortex and reactive gliosis in the brain. Genes Dev. 2001;15(7):859–76.

20. Gregorian C, Nakashima J, Le Belle J, Ohab J, Kim R, Liu A, et al. Pten Deletion in Adult Neural Stem/Progenitor Cells Enhances Constitutive Neurogenesis. The Journal of Neuroscience. 2009;29(6):1874.

21. Reekes TH, Higginson CI, Ledbetter CR, Sathivadivel N, Zweig RM, Disbrow EA. Sex specific cognitive differences in Parkinson disease. npj Parkinson’s Disease. 2020;6(1):7.

22. Oakley H, Cole SL, Logan S, Maus E, Shao P, Craft J, et al. Intraneuronal β-Amyloid Aggregates, Neurodegeneration, and Neuron Loss in Transgenic Mice with Five Familial Alzheimer’s Disease Mutations: Potential Factors in Amyloid Plaque Formation. The Journal of Neuroscience. 2006;26(40):10129.

23. Goedert M, Spillantini MG, Del Tredici K, Braak H. 100 years of Lewy pathology. Nature Reviews Neurology. 2013;9(1):13–24.

24. Zarei M, Ibarretxe-Bilbao N, Compta Y, Hough M, Junque C, Bargallo N, et al. Cortical thinning is associated with disease stages and dementia in Parkinson&#039;s disease. Journal of Neurology, Neurosurgery & Psychiatry. 2013;84(8):875.

25. Compta Y, Parkkinen L, Kempster P, Selikhova M, Lashley T, Holton JL, et al. The Significance of α-Synuclein, Amyloid-β and Tau Pathologies in Parkinson’s Disease Progression and Related Dementia. Neurodegenerative Diseases. 2014;13(2-3):154–6.

26. Irwin DJ, Lee VMY, Trojanowski JQ. Parkinson’s disease dementia: convergence of α-synuclein, tau and amyloid-β pathologies. Nature Reviews Neuroscience. 2013;14:626.

27. Camicioli R, Moore MM, Kinney A, Corbridge E, Glassberg K, Kaye JA. Parkinson’s disease is associated with hippocampal atrophy. Movement Disorders. 2003;18(7):784–90.

28. Doty RL. Olfactory dysfunction in Parkinson disease. Nature Reviews Neurology. 2012;8(6):329–39.

29. Pfisterer U, Petukhov V, Demharter S, Meichsner J, Thompson JJ, Batiuk MY, et al. Identification of epilepsy-associated neuronal subtypes and gene expression underlying epileptogenesis. Nature Communications. 2020;11(1):5038.

30. Krishnaswami SR, Grindberg RV, Novotny M, Venepally P, Lacar B, Bhutani K, et al. Using single nuclei for RNA-seq to capture the transcriptome of postmortem neurons. Nature Protocols. 2016;11(3):499–524.

31. Stuart T, Butler A, Hoffman P, Hafemeister C, Papalexi E, Mauck WM, et al. Comprehensive Integration of Single-Cell Data. Cell. 2019;177(7):1888–902.e21.

32. Barkas N, Petukhov V, Nikolaeva D, Lozinsky Y, Demharter S, Khodosevich K, et al. Joint analysis of heterogeneous single-cell RNA-seq dataset collections. Nature Methods. 2019;16(8):695–8.

33. Wolock SL, Lopez R, Klein AM. Scrublet: Computational Identification of Cell Doublets in Single-Cell Transcriptomic Data. Cell systems. 2019;8(4):281–91.e9.

34. Haghverdi L, Lun ATL, Morgan MD, Marioni JC. Batch effects in single-cell RNA-sequencing data are corrected by matching mutual nearest neighbors. Nature Biotechnology. 2018;36(5):421–7.

35. Subramanian A, Tamayo P, Mootha VK, Mukherjee S, Ebert BL, Gillette MA, et al. Gene set enrichment analysis: A knowledge-based approach for interpreting genome-wide expression profiles. Proceedings of the National Academy of Sciences. 2005;102(43):15545.

36. Liberzon A, Birger C, Thorvaldsdóttir H, Ghandi M, Mesirov Jill P, Tamayo P. The Molecular Signatures Database Hallmark Gene Set Collection. Cell systems. 2015;1(6):417–25.

37. Heberle H, Meirelles GV, da Silva FR, Telles GP, Minghim R. InteractiVenn: a web-based tool for the analysis of sets through Venn diagrams. BMC Bioinformatics. 2015;16(1):169.

38. Kulak NA, Geyer PE, Mann M. Loss-less Nano-fractionator for High Sensitivity, High Coverage Proteomics. Molecular & cellular proteomics : MCP. 2017;16(4):694–705.

39. Brunner A-D, Thielert M, Vasilopoulou CG, Ammar C, Coscia F, Mund A, et al. Ultra-high sensitivity mass spectrometry quantifies single-cell proteome changes upon perturbation. bioRxiv. 2021:2020.12.22.423933.

40. Santos A, Colaço AR, Nielsen AB, Niu L, Geyer PE, Coscia F, et al. Clinical Knowledge Graph Integrates Proteomics Data into Clinical Decision-Making. bioRxiv. 2020:2020.05.09.084897.

41. Andersen JV, Skotte NH, Christensen SK, Polli FS, Shabani M, Markussen KH, et al. Hippocampal disruptions of synaptic and astrocyte metabolism are primary events of early amyloid pathology in the 5xFAD mouse model of Alzheimer’s disease. Cell Death & Disease. 2021;12(11):954.

42. Skotte NH, Andersen JV, Santos A, Aldana BI, Willert CW, Nørremølle A, et al. Integrative Characterization of the R6/2 Mouse Model of Huntington’s Disease Reveals Dysfunctional Astrocyte Metabolism. Cell reports. 2018;23(7):2211–24.

43. Andersen JV, Jakobsen E, Westi EW, Lie MEK, Voss CM, Aldana BI, et al. Extensive astrocyte metabolism of γ-aminobutyric acid (GABA) sustains glutamine synthesis in the mammalian cerebral cortex. Glia. 2020.

44. Westi EW, Andersen JV, Aldana BI. Using stable isotope tracing to unravel the metabolic components of neurodegeneration: Focus on neuron-glia metabolic interactions. Neurobiol Dis. 2023;182:106145.

45. Southwell AL, Warby SC, Carroll JB, Doty CN, Skotte NH, Zhang W, et al. A fully humanized transgenic mouse model of Huntington disease. Human molecular genetics. 2013;22(1):18–34.

46. Samuels BA, Hen R. Novelty-Suppressed Feeding in the Mouse. In: Gould TD, editor. Mood and Anxiety Related Phenotypes in Mice: Characterization Using Behavioral Tests, Volume II. Totowa, NJ: Humana Press; 2011. p. 107–21.

47. Santarelli L, Saxe M, Gross C, Surget A, Battaglia F, Dulawa S, et al. Requirement of Hippocampal Neurogenesis for the Behavioral Effects of Antidepressants. Science. 2003;301(5634):805-9.

48. Liu M-Y, Yin C-Y, Zhu L-J, Zhu X-H, Xu C, Luo C-X, et al. Sucrose preference test for measurement of stress-induced anhedonia in mice. Nature Protocols. 2018;13(7):1686–98.

49. Southwell AL, Ko J, Patterson PH. Intrabody gene therapy ameliorates motor, cognitive, and neuropathological symptoms in multiple mouse models of Huntington’s disease. The Journal of neuroscience : the official journal of the Society for Neuroscience. 2009;29(43):13589–602.

50. Liu Y, Marin A, Ejlerskov P, Rasmussen LM, Prinz M, Issazadeh-Navikas S. Neuronal IFN-beta-induced PI3K/Akt-FoxA1 signalling is essential for generation of FoxA1+Treg cells. Nat Commun. 2017;8:14709.

51. Blank T, Detje CN, Spiess A, Hagemeyer N, Brendecke SM, Wolfart J, et al. Brain Endothelial- and Epithelial-Specific Interferon Receptor Chain 1 Drives Virus-Induced Sickness Behavior and Cognitive Impairment. Immunity. 2016;44(4):901–12.

52. Tresse E, Riera-Ponsati L, Jaberi E, Sew WQG, Ruscher K, Issazadeh-Navikas S. IFN-β rescues neurodegeneration by regulating mitochondrial fission via STAT5, PGAM5, and Drp1. The EMBO Journal. 2021;n/a(n/a):e106868.

53. Kwon D, Kim C, Woo YK, Hwang JK. Inhibitory Effects of Chrysanthemum (Chrysanthemum morifolium Ramat.) Extract and Its Active Compound Isochlorogenic Acid A on Sarcopenia. Prev Nutr Food Sci. 2021;26(4):408–16.

54. Burmeister AR, Johnson MB, Marriott I. Murine astrocytes are responsive to the pro-inflammatory effects of IL-20. Neurosci Lett. 2019;708:134334.

55. Jana A, Wang X, Leasure JW, Magana L, Wang L, Kim YM, et al. Increased Type I interferon signaling and brain endothelial barrier dysfunction in an experimental model of Alzheimer’s disease. Scientific reports. 2022;12(1):16488.

56. Perng Y-C, Lenschow DJ. ISG15 in antiviral immunity and beyond. Nature Reviews Microbiology. 2018;16(7):423–39.

57. Castiglia V, Piersigilli A, Ebner F, Janos M, Goldmann O, Damböck U, et al. Type I Interferon Signaling Prevents IL-1β-Driven Lethal Systemic Hyperinflammation during Invasive Bacterial Infection of Soft Tissue. Cell Host & Microbe. 2016;19(3):375–87.

58. Lester DB, Rogers TD, Blaha CD. Acetylcholine–Dopamine Interactions in the Pathophysiology and Treatment of CNS Disorders. CNS Neuroscience & Therapeutics. 2010;16(3):137–62.

59. Born G, Grayton HM, Langhorst H, Dudanova I, Rohlmann A, Woodward BW, et al. Genetic targeting of NRXN2 in mice unveils role in excitatory cortical synapse function and social behaviors. Front Synaptic Neurosci. 2015;7:3-.

60. Bandres-Ciga S, Saez-Atienzar S, Bonet-Ponce L, Billingsley K, Vitale D, Blauwendraat C, et al. The endocytic membrane trafficking pathway plays a major role in the risk of Parkinson’s disease. Movement Disorders. 2019;34(4):460–8.

61. Hondius DC, Koopmans F, Leistner C, Pita-Illobre D, Peferoen-Baert RM, Marbus F, et al. The proteome of granulovacuolar degeneration and neurofibrillary tangles in Alzheimer’s disease. Acta Neuropathologica. 2021;141(3):341–58.

62. Miyashita A, Hatsuta H, Kikuchi M, Nakaya A, Saito Y, Tsukie T, et al. Genes associated with the progression of neurofibrillary tangles in Alzheimer’s disease. Translational Psychiatry. 2014;4(6):e396-e.

63. Parcellier A, Tintignac LA, Zhuravleva E, Dummler B, Brazil DP, Hynx D, et al. The Carboxy-Terminal Modulator Protein (CTMP) regulates mitochondrial dynamics. PLoS One. 2009;4(5):e5471.

64. Parcellier A, Tintignac LA, Zhuravleva E, Cron P, Schenk S, Bozulic L, et al. Carboxy-Terminal Modulator Protein (CTMP) is a mitochondrial protein that sensitizes cells to apoptosis. Cell Signal. 2009;21(4):639–50.

65. Ojiakor OA, Rylett RJ. Modulation of sodium-coupled choline transporter CHT function in health and disease. Neurochemistry International. 2020;140:104810.

66. Jaudon F, Raynaud F, Wehrlé R, Bellanger J-M, Doulazmi M, Vodjdani G, et al. The RhoGEF DOCK10 is essential for dendritic spine morphogenesis. Mol Biol Cell. 2015;26(11):2112–27.

67. Wong YC, Holzbaur EL. Optineurin is an autophagy receptor for damaged mitochondria in parkin-mediated mitophagy that is disrupted by an ALS-linked mutation. Proc Natl Acad Sci U S A. 2014;111(42):E4439–48.

68. Imberechts D, Kinnart I, Wauters F, Terbeek J, Manders L, Wierda K, et al. DJ-1 is an essential downstream mediator in PINK1/parkin-dependent mitophagy. Brain. 2022;145(12):4368–84.

69. Andersen JV, Schousboe A. Milestone Review: Metabolic dynamics of glutamate and GABA mediated neurotransmission - The essential roles of astrocytes. J Neurochem. 2023;166(2):109–37.

70. Wyss MT, Magistretti PJ, Buck A, Weber B. Labeled acetate as a marker of astrocytic metabolism. J Cereb Blood Flow Metab. 2011;31(8):1668–74.

71. Alberico SL, Cassell MD, Narayanan NS. The Vulnerable Ventral Tegmental Area in Parkinson’s Disease. Basal Ganglia. 2015;5(2-3):51–5.

72. Jacob EL, Gatto NM, Thompson A, Bordelon Y, Ritz B. Occurrence of depression and anxiety prior to Parkinson’s disease. Parkinsonism & related disorders. 2010;16(9):576–81.

73. Lin CH, Wu RM, Chang HY, Chiang YT, Lin HH. Preceding pain symptoms and Parkinson’s disease: a nationwide population-based cohort study. European Journal of Neurology. 2013;20(10):1398–404.

74. Oaks AW, Frankfurt M, Finkelstein DI, Sidhu A. Age-Dependent Effects of A53T Alpha-Synuclein on Behavior and Dopaminergic Function. PLOS ONE. 2013;8(4):e60378.

75. Branchi I, D’Andrea I, Armida M, Carnevale D, Ajmone-Cat MA, Pèzzola A, et al. Striatal 6-OHDA lesion in mice: Investigating early neurochemical changes underlying Parkinson’s disease. Behav Brain Res. 2010;208(1):137–43.

76. Murray C, Griffin É W, O’Loughlin E, Lyons A, Sherwin E, Ahmed S, et al. Interdependent and independent roles of type I interferons and IL-6 in innate immune, neuroinflammatory and sickness behaviour responses to systemic poly I:C. Brain, behavior, and immunity. 2015;48:274–86.

77. Mitani Y, Takaoka A, Kim SH, Kato Y, Yokochi T, Tanaka N, et al. Cross talk of the interferon-alpha/beta signalling complex with gp130 for effective interleukin-6 signalling. Genes Cells. 2001;6(7):631–40.

78. He X-f, Xu J-h, Li G, Li M-y, Li L-l, Pei Z, et al. NLRP3-dependent microglial training impaired the clearance of amyloid-beta and aggravated the cognitive decline in Alzheimer’s disease. Cell Death & Disease. 2020;11(10):849.

79. Zenaro E, Piacentino G, Constantin G. The blood-brain barrier in Alzheimer’s disease. Neurobiology of Disease. 2017;107:41–56.

80. Raz L, Knoefel J, Bhaskar K. The neuropathology and cerebrovascular mechanisms of dementia. J Cereb Blood Flow Metab. 2016;36(1):172–86.

81. Su M, Hu H, Lee Y, D’Azzo A, Messing A, Brenner M. Expression Specificity of GFAP Transgenes. Neurochemical research. 2004;29(11):2075–93.

82. Srinivasan R, Lu T-Y, Chai H, Xu J, Huang BS, Golshani P, et al. New Transgenic Mouse Lines for Selectively Targeting Astrocytes and Studying Calcium Signals in Astrocyte Processes In Situ and In Vivo. Neuron. 2016;92(6):1181–95.

83. Giusti SA, Vercelli CA, Vogl AM, Kolarz AW, Pino NS, Deussing JM, et al. Behavioral phenotyping of Nestin-Cre mice: Implications for genetic mouse models of psychiatric disorders. Journal of Psychiatric Research. 2014;55:87–95.

84. Sirkis DW, Oddi AP, Jonson C, Bonham LW, Hoang PT, Yokoyama JS. The role of interferon signaling in neurodegeneration and neuropsychiatric disorders. Front Psychiatry. 2024;15:1480438.

85. Chhatbar C, Detje CN, Grabski E, Borst K, Spanier J, Ghita L, et al. Type I Interferon Receptor Signaling of Neurons and Astrocytes Regulates Microglia Activation during Viral Encephalitis. Cell reports. 2018;25(1):118–29.e4.

86. Hayes CK, Giraldo D, Wilcox DR, Longnecker R. The Astrocyte Type I Interferon Response Is Essential for Protection against Herpes Simplex Encephalitis. J Virol. 2022;96(4):e0178321.

87. Chotiwan N, Rosendal E, Willekens SMA, Schexnaydre E, Nilsson E, Lindqvist R, et al. Type I interferon shapes brain distribution and tropism of tick-borne flavivirus. Nature Communications. 2023;14(1):2007.

88. Nehammer C, Ejlerskov P, Gopal S, Handley A, Ng L, Moreira P, et al. Interferon-β-induced miR-1 alleviates toxic protein accumulation by controlling autophagy. eLife. 2019;8:e49930.

89. Gatt AP, Duncan OF, Attems J, Francis PT, Ballard CG, Bateman JM. Dementia in Parkinson’s disease is associated with enhanced mitochondrial complex I deficiency. Mov Disord. 2016;31(3):352–9.

90. Garcia-Esparcia P, Koneti A, Rodríguez-Oroz MC, Gago B, Del Rio JA, Ferrer I. Mitochondrial activity in the frontal cortex area 8 and angular gyrus in Parkinson’s disease and Parkinson’s disease with dementia. Brain pathology (Zurich, Switzerland). 2018;28(1):43–57.

91. Wu D, Sanin DE, Everts B, Chen Q, Qiu J, Buck MD, et al. Type 1 Interferons Induce Changes in Core Metabolism that Are Critical for Immune Function. Immunity. 2016;44(6):1325–36.

92. Burke JD, Platanias LC, Fish EN. Beta Interferon Regulation of Glucose Metabolism Is PI3K/Akt Dependent and Important for Antiviral Activity against Coxsackievirus B3. Journal of Virology. 2014;88(6):3485.

93. Sibson NR, Dhankhar A, Mason GF, Rothman DL, Behar KL, Shulman RG. Stoichiometric coupling of brain glucose metabolism and glutamatergic neuronal activity. Proceedings of the National Academy of Sciences. 1998;95(1):316.

94. Nilsen LH, Rae C, Ittner LM, Götz J, Sonnewald U. Glutamate metabolism is impaired in transgenic mice with tau hyperphosphorylation. J Cereb Blood Flow Metab. 2013;33(5):684–91.

95. Ibarretxe-Bilbao N, Ramírez-Ruiz B, Tolosa E, Martí MJ, Valldeoriola F, Bargalló N, et al. Hippocampal head atrophy predominance in Parkinson’s disease with hallucinations and with dementia. Journal of Neurology. 2008;255(9):1324–31.

96. Dickson DW, Braak H, Duda JE, Duyckaerts C, Gasser T, Halliday GM, et al. Neuropathological assessment of Parkinson’s disease: refining the diagnostic criteria. The Lancet Neurology. 2009;8(12):1150–7.

97. Molloy SA, Rowan EN, O’Brien JT, McKeith IG, Wesnes K, Burn DJ. Effect of levodopa on cognitive function in Parkinson’s disease with and without dementia and dementia with Lewy bodies. J Neurol Neurosurg Psychiatry. 2006;77(12):1323–8.

98. Fabbri M, Coelho M, Guedes LC, Chendo I, Sousa C, Rosa MM, et al. Response of non-motor symptoms to levodopa in late-stage Parkinson’s disease: Results of a levodopa challenge test. Parkinsonism Relat Disord. 2017;39:37–43.

99. Williams L, Qiu J, Waller S, Tsui D, Griffith J, Fung VSC. Challenges in managing late-stage Parkinson’s disease: Practical approaches and pitfalls. Aust J Gen Pract. 2022;51(10):778–85.

100. Lin WY, Lin MS, Weng YH, Yeh TH, Lin YS, Fong PY, et al. Association of Antiviral Therapy With Risk of Parkinson Disease in Patients With Chronic Hepatitis C Virus Infection. JAMA Neurol. 2019;76(9):1019–27.

101. Liu Y, Carlsson R, Ambjorn M, Hasan M, Badn W, Darabi A, et al. PD-L1 expression by neurons nearby tumors indicates better prognosis in glioblastoma patients. J Neurosci. 2013;33(35):14231–45.

102. de Weerd NA, Vivian JP, Nguyen TK, Mangan NE, Gould JA, Braniff S-J, et al. Structural basis of a unique interferon-β signaling axis mediated via the receptor IFNAR1. Nat Immunol. 2013;14(9):901–7.

103. Khakh BS. Astrocyte-Neuron Interactions in the Striatum: Insights on Identity, Form, and Function. Trends Neurosci. 2019;42(9):617–30.

104. Chai H, Diaz-Castro B, Shigetomi E, Monte E, Octeau JC, Yu X, et al. Neural Circuit-Specialized Astrocytes: Transcriptomic, Proteomic, Morphological, and Functional Evidence. Neuron. 2017;95(3):531–49.e9.

105. Perez-Riverol Y, Csordas A, Bai J, Bernal-Llinares M, Hewapathirana S, Kundu DJ, et al. The PRIDE database and related tools and resources in 2019: improving support for quantification data. Nucleic acids research. 2019;47(D1):D442–D50.

