## Supplementary figures and images for "Distinct and Combined Interferon-ɑ/β-receptor-1 Loss in Neurons and Astrocytes Disrupt Brain Energy Metabolism and Drive Parkinsonian Dementia"

### Supplementary Fig. 1

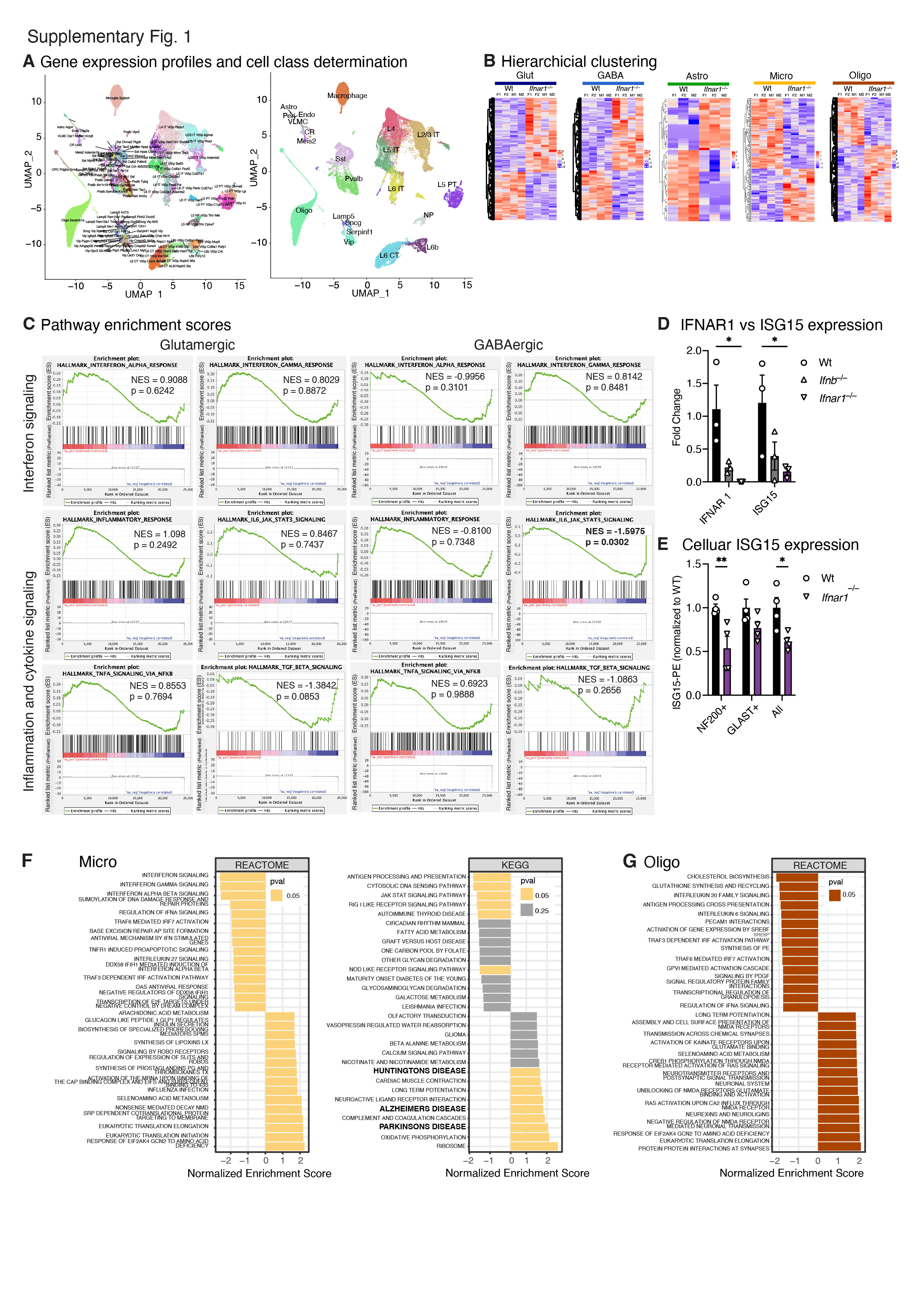

### Supplementary Fig. 2

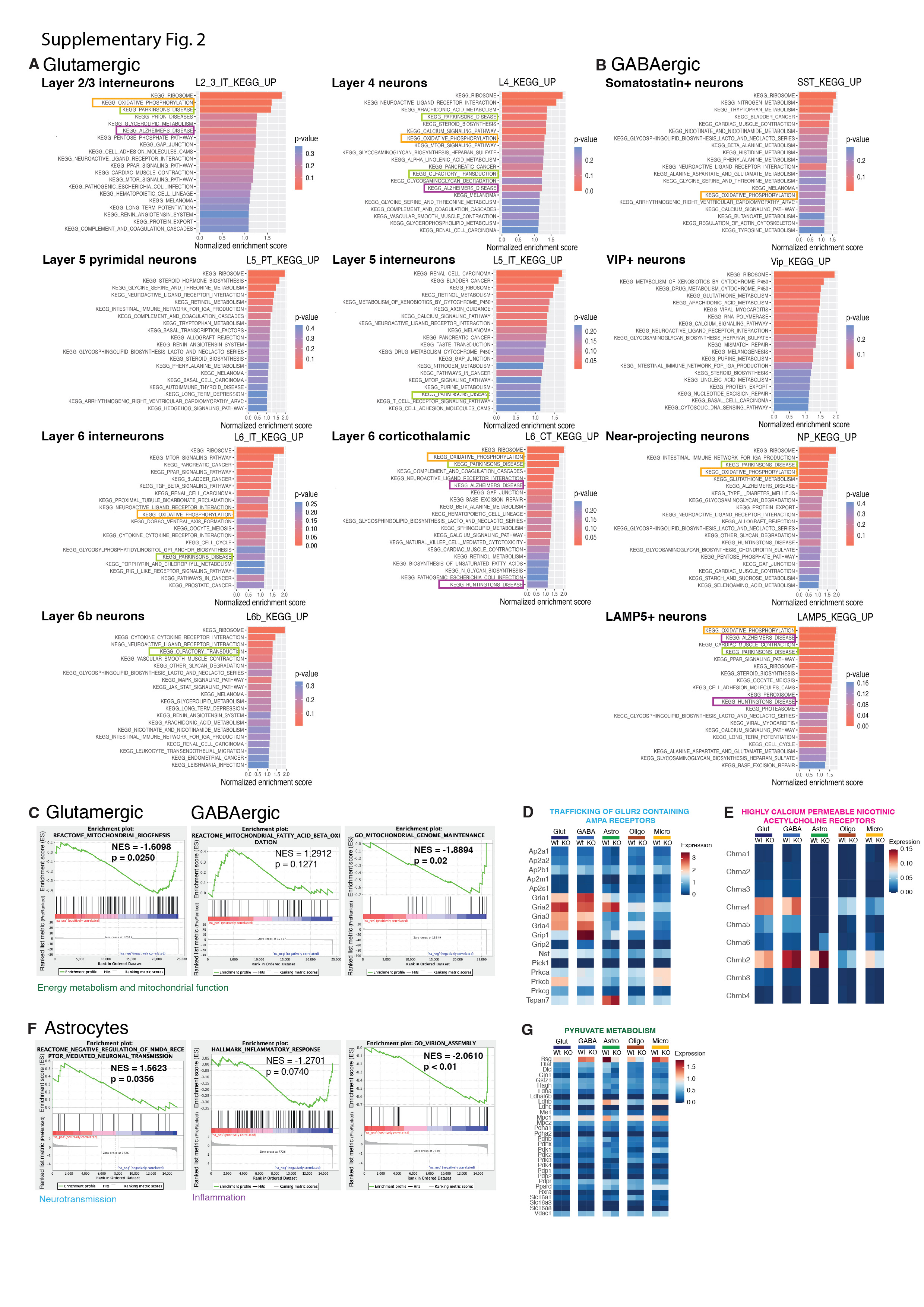

### Supplementary Fig. 3

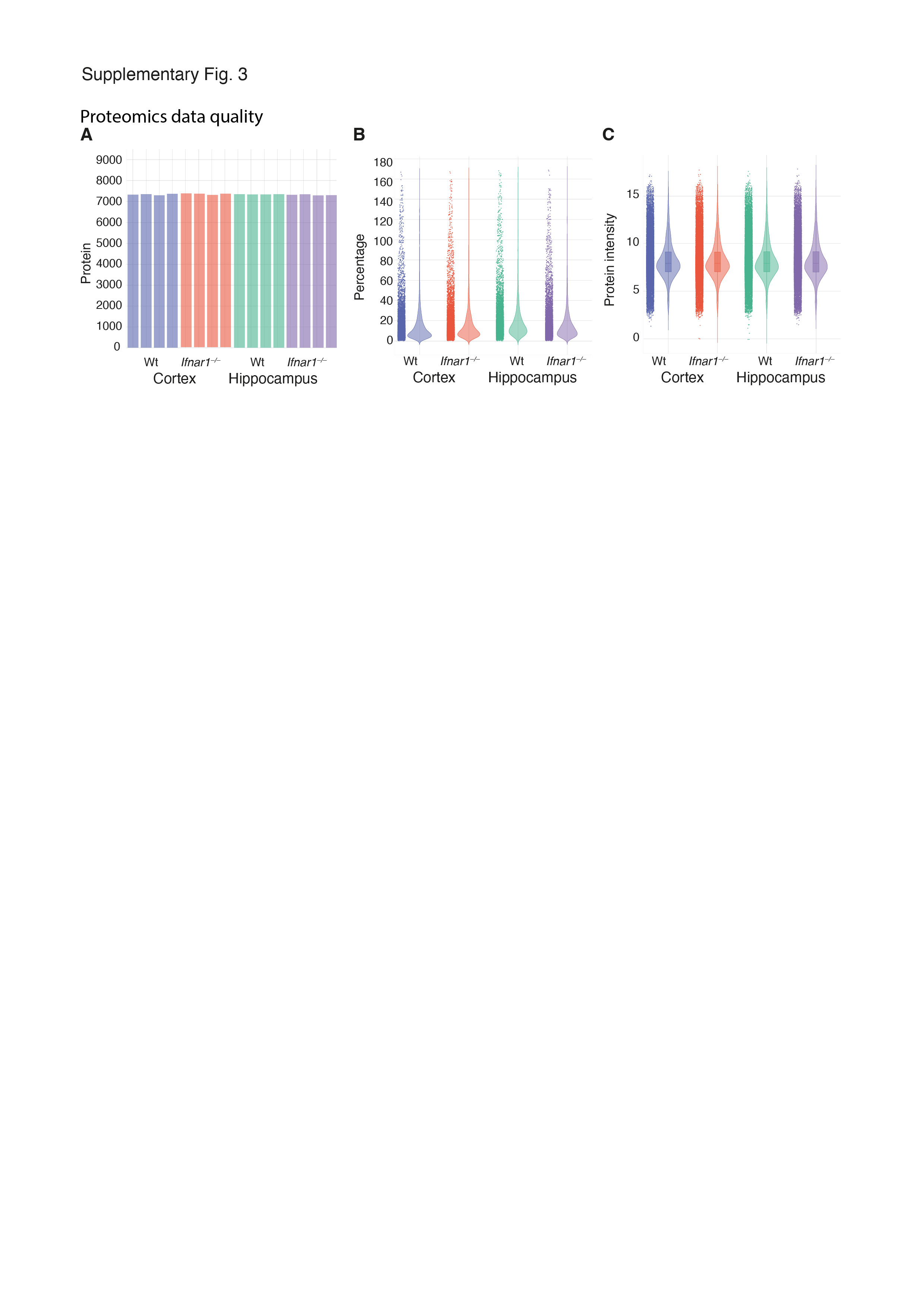

### Supplementary Fig. 4

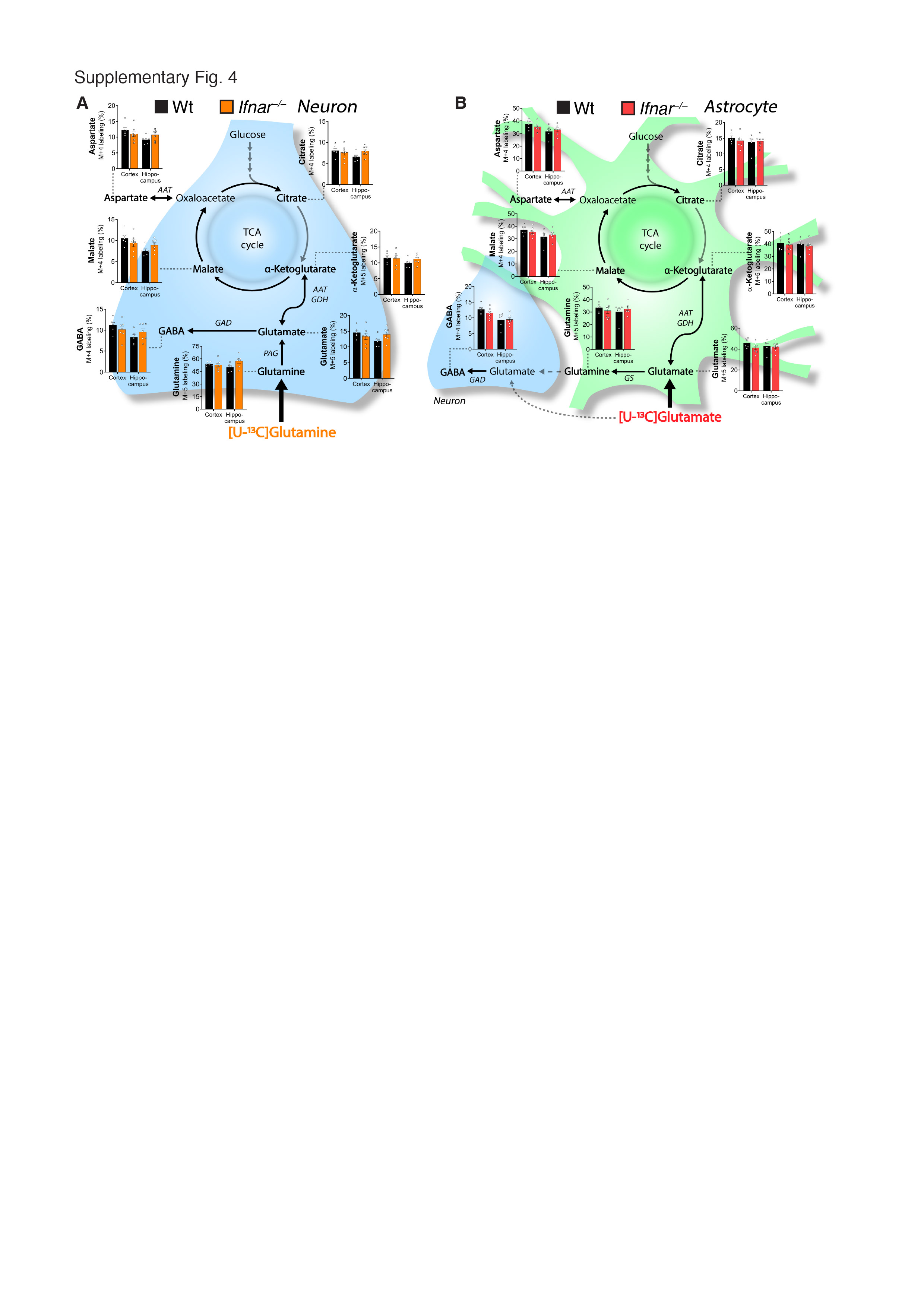

### Supplementary Fig. 5

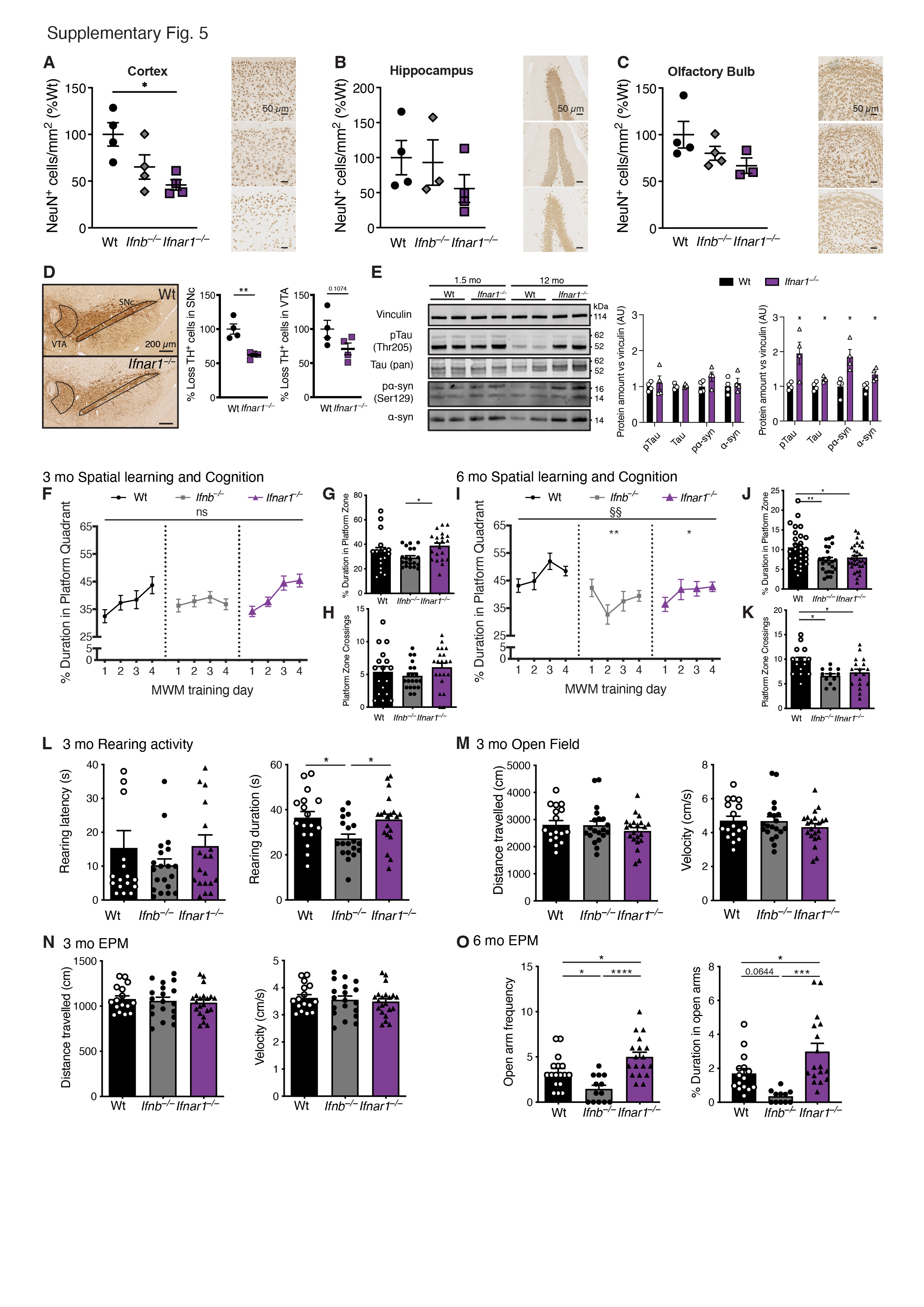

### Supplementary Fig. 6

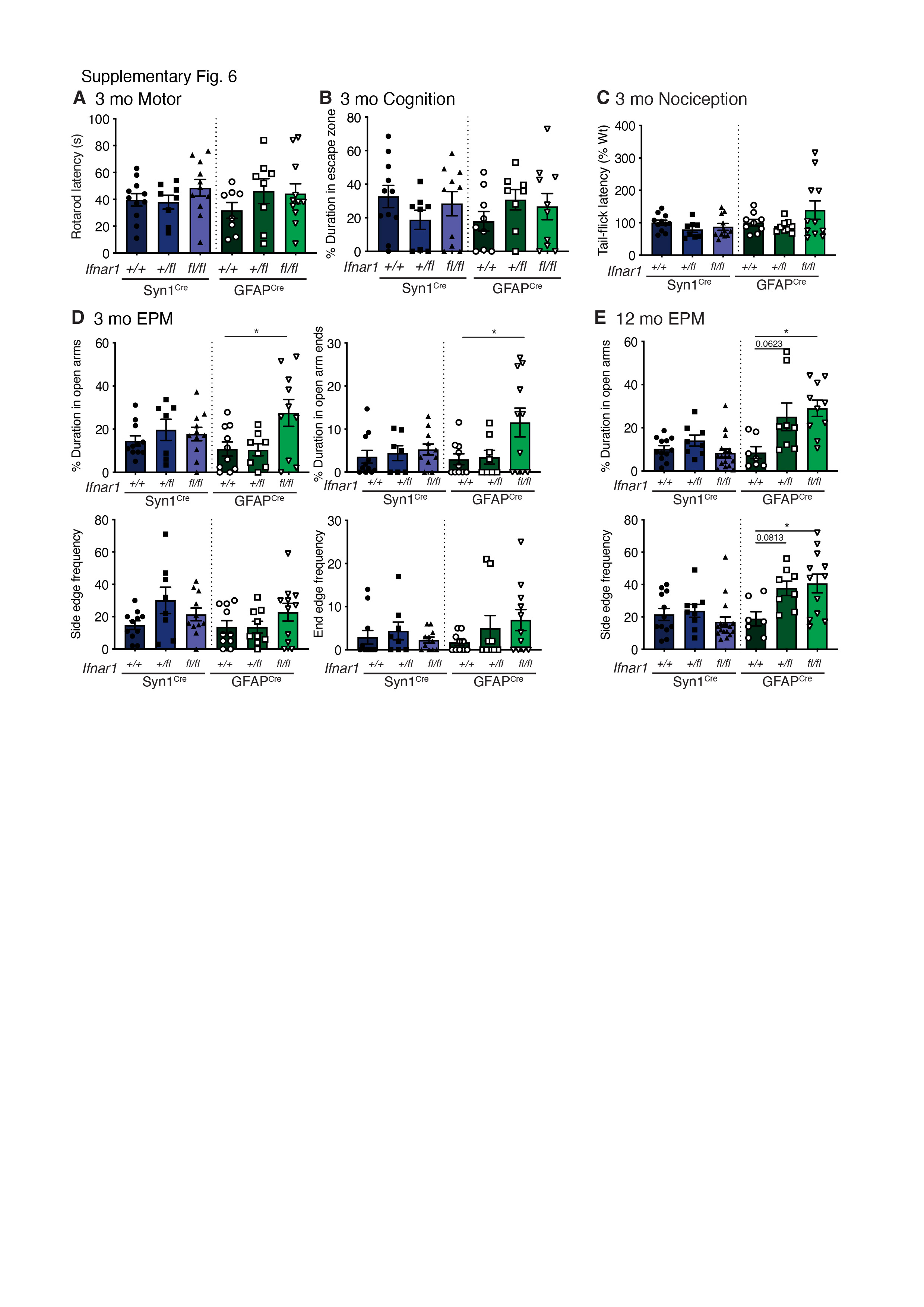
